# Dietary methionine restriction impairs anti-tumor immunity through gut microbiota

**DOI:** 10.1101/2021.08.27.457955

**Authors:** Ming Ji, Xiaojiang Xu, Qing Xu, Xin Xu, M. Andrea Azcarate-Peril, Xiaoyue Wu, Juan Liu, Jason W. Locasale, Jian-Liang Li, Igor Shats, Xiaoling Li

**Author notes:** Correspondence (X. L.).

## Abstract

Dietary methionine restriction has been reported to repress cancer growth and improve therapeutic responses in several pre-clinical settings. However, how this dietary intervention impacts cancer progression in the context of the intact immune system is unknown. Here we report that methionine restriction exacerbates cancer growth and influences the outcomes of anti-tumor immunotherapy through gut microbiota and immune suppression in immunocompetent settings. Methionine restriction reduces T cell activation, increases tumor growth, and impairs response to anti-tumor immunotherapy. Mechanistically, methionine restriction alters composition of gut microbiota and reduces microbial production of hydrogen sulfide. Fecal transplantation from methionine-restricted tumor-free animals is sufficient to repress T cell activation and enhance tumor growth in tumor-bearing recipient mice. Conversely, dietary supplementation of a hydrogen sulfide donor or methionine stimulates anti-tumor immunity and suppresses tumor progression. Our findings reveal a vital role of gut microbiota in mediating methionine restriction-induced suppression of anti-tumor immunity and suggest that any possible anti-cancer benefits of methionine restriction require careful considerations of both the microbiota and the immune system.

## Introduction

Methionine, a sulfur-containing essential amino acid, is a key component of dietary proteins important for protein synthesis, sulfur metabolism, epigenetic modification, antioxidant defense, and signaling (Mato et al., 2008). However, the role of methionine in regulation of cancer progression remains inconclusive. On one hand, many cancer cells, including tumor-initiating cells and HNF4*α*-positive liver cancer cells, are highly dependent on exogeneous methionine (Wang et al., 2019; Xu et al., 2020). Methionine restriction (MR), a dietary regimen that protects against metabolic diseases and aging (Ables et al., 2016; Ables and Johnson, 2017; Miller et al., 2005; Orentreich et al., 1993; Sun et al., 2009), is known to repress proliferation and progression of a variety of xenograft tumors (Ables and Johnson, 2017; Guo et al., 1993; Hens et al., 2016; Komninou et al., 2006; Poirson-Bichat et al., 1997; Sinha et al., 2014; Xu et al., 2020). This dietary intervention can impact metabolic flux in one-carbon metabolism, inhibit tumor growth, and sensitize tumors to chemotherapy and radiotherapy in certain patient-derived xenografts (PDXs) in a tumor-cell autonomous manner (Gao et al., 2019). On the other hand, methionine is also critical for T cell activation and differentiation (Roy et al., 2020; Sinclair et al., 2019). Competition for methionine between fast-growing tumor cells and T cells in the tumor microenvironment has been reported to disrupt methionine metabolism inside CD8^+^ T cells, lowering intracellular levels of methionine and the methyl donor S-adenosylmethionine (SAM) (Bian et al., 2020). This reduction leads to loss of H3K79me2 histone mark, decreased expression of STAT5, and impairment of T cell immunity and checkpoint-induced tumor immunity (Bian et al., 2020). Therefore, methionine is also a potential tumor suppressing nutrient that enhances T cell-mediated anti-tumor immunity.

In this study, we investigated the interaction between dietary methionine, immune cells, and cancer cells in immunocompetent mice. Using genetic and allograft tumor models, together with transcriptomic profiling, scRNA-seq analysis, 16S rRNA gene amplicon sequencing, and fecal transplantation, we uncover an intriguing mechanism in which gut microbiota couples dietary methionine to T cell activation and anti-tumor immunity in immunocompetent mice.

## Results

### Dietary methionine restriction enhances tumor progression and represses T cell activation in *Apc^min+/-^* mice

To investigate the potential impact of dietary methionine restriction on intestinal tumor progression, we fed a genetic intestinal tumor model, *Apc^min+/-^* mice (Moser et al., 1990; Su et al., 1992), with either a methionine-restricted diet (MR diet) containing 0.172% DL-methionine or a control diet (CTRL diet) containing 0.86% DL-methionine at the age of eight weeks, and analyzed the effects of these diets on intestinal tumor growth eight weeks later. Since a number of comparable methionine-restricted diets have been shown to suppress a variety of xenograft tumors (Ables and Johnson, 2017; Guo et al., 1993; Hens et al., 2016; Komninou et al., 2006; Poirson-Bichat et al., 1997; Sinha et al., 2014), including colorectal PDX tumors in mice (Gao et al., 2019), we were surprised to find that *Apc^min+/-^* mice fed with the MR diet had a significantly higher number of tumors in the small intestine and a higher tumor burden in the colon compared to those fed with the control diet (Figure 1A and 1B). Specifically, *Apc^min+/-^* mice on the MR diet developed more polyps in the ileum and more adenomas in the colon than *Apc^min+/-^* mice on the control diet (Figure 1C). Moreover, the increase of tumor number in the ileum of the MR diet fed mice was coupled with reduced expression levels of several apoptotic and DNA damage response genes (Figure S1A), decreased cleavage of apoptotic protein Bax (Figure S1B), and an increased trend of Ki67-positive staining, a marker for cell proliferation (Figure 1D), indicating that intestinal tumors developed on the MR diet are more resistant to cell death and may be more proliferative than intestinal tumors developed on the control diet.

**Figure 1.**
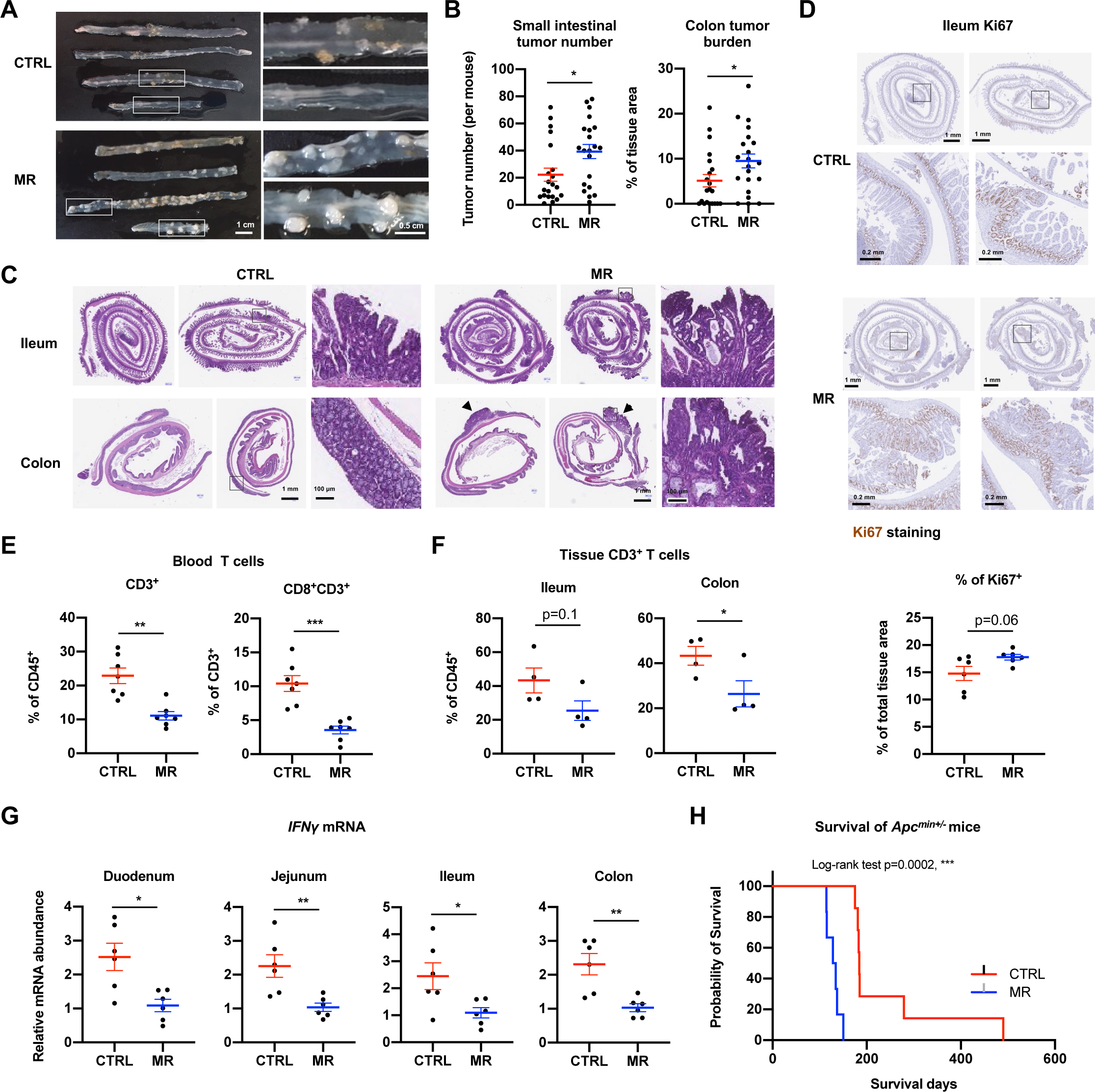
Dietary methionine restriction enhances tumor progression and represses T cell activation in *Apc^min+/-^* mice. (A-C) Dietary methionine restriction increases intestinal tumor growth in *Apc^min+/-^* mice. *Apc^min+/-^* mice were fed with a control (CTRL) diet containing 0.86% of methionine and a methionine restricted (MR) diet containing 0.172% methionine as described in Methods. (A) Representative gross morphological images of intestinal tumors. (B) Small intestinal tumor numbers and colonic tumor burden. The final tumor number in the small intestine and the tumor burden in the colon were calculated as described in Methods (values are expressed as mean ± s.e.m., n=21 mice in each group, *p<0.05). (C) Representative images of H&E-stained ileum and colon sections. (D) Dietary methionine restriction increases epithelial cell proliferation in the ileum of *Apc^min+/-^* mice. The ileum sections from mice fed with control diet and methionine restricted diet were stained by Ki67 as described in Methods. Representative images are shown. (E) Dietary methionine restriction reduces the fraction of circulating CD3^+^ T cells in *Apc^min+/-^* mice. The abundance of CD3^+^ and CD3^+^ CD8^+^ T cells in the blood of *Apc^min+/-^* mice on CTRL diet or MR diet was analyzed by flow cytometry as described in Methods (values are expressed as mean ± s.e.m., n=7 mice in each group, ***p<0.001, **p<0.01). (F) Dietary methionine restriction reduces the fraction of CD3^+^ T cells in intestinal tissues of *Apc^min+/-^* mice (values are expressed as mean ± s.e.m., n=3 or 4 mice in each group, *p<0.05). (G) Dietary methionine restriction reduces the expression of *IFNγ* in all segments of the intestine. The mRNA levels of *IFNγ* were analyzed by qPCR (values are expressed as mean ± s.e.m., n=6 mice in each group, ***p<0.001, **p<0.01). (H) Dietary methionine restriction reduces survival of *Apc^min+/-^* mice. The survival of *Apc^min+/-^* mice under the indicated diets were monitored (n=7 mice on CTRL diet and 6 mice on MR diet).

*Apc^min+/-^* mice are immunocompetent in contrast with immunodeficient mice utilized in previous xenograft and PDX studies (Ables and Johnson, 2017; Gao et al., 2019; Guo et al., 1993; Hens et al., 2016; Komninou et al., 2006; Poirson-Bichat et al., 1997; Sinha et al., 2014). To comprehend this unexpected tumor-promoting effect of dietary methionine restriction in *Apc^min+/-^* mice, we tested whether dietary methionine restriction impaired T cell differentiation and activation, which are known to be supported by methionine (Bian et al., 2020; Roy et al., 2020; Sinclair et al., 2019). When quantified by flow cytometry, MR diet feeding significantly reduced the fraction of circulating CD3^+^ and CD8^+^CD3^+^ T cells in the blood (Figure 1E and S1C), along with decreased abundance of CD3^+^ T cells in the ileum and colon (Figure 1F). In line with these observations, the mRNA levels of *IFNγ,* a key tumor immune surveillance cytokine predominantly produced by activated lymphocytes, including CD4^+^ T helper type 1 (Th1) cells, CD8^+^ cytotoxic T cells, natural killer T (NKT) cells, natural killer cells (NK), and some innate lymphoid cells (ILCs) (Castro et al., 2018), were significantly repressed in all segments of the intestine of MR diet-fed *Apc^min+/-^* mice (Figure 1G). These observations indicate that dietary methionine restriction-induced tumor growth in immunocompetent *Apc^min+/-^* mice is associated with impaired immune function. Notably, the median health-span (symptom-free survival) of *Apc^min+/-^* mice under the control diet was 184 days. Dietary methionine restriction dramatically shortened their median health-span to 131 days (Figure 1H). Taken together, our observations indicate that dietary methionine restriction is associated with accelerated tumor progression and reduced T cell population in immunocompetent *Apc^min+/-^* mice.

### Dietary methionine restriction represses immune activation and reduces efficacy of anti-tumor immunotherapy in immunocompetent mice

The increase in intestinal tumor burden and reduction of circulating and tissue CD3^+^ T cells in MR diet-fed *Apc^min+/-^* mice raise the possibility that dietary methionine restriction may promote tumor progression by suppressing immune activation and anti-tumor immunity in immunocompetent mice. To directly evaluate the role of the immune system in the response of tumors to dietary methionine restriction, we compared the effects of MR diet feeding on the growth of allograft CT26.WT mouse colon carcinoma cells in immunocompetent Balb/c mice and immunocompromised NSG mice, one of the most immunocompromised mouse strains that lacks mature T, B, and natural killer cells due to the *scid* mutation and *Il2rg* knockout (Shultz et al., 2005). Consistent with previous reports (Ables and Johnson, 2017; Guo et al., 1993; Hens et al., 2016; Komninou et al., 2006; Poirson-Bichat et al., 1997; Sinha et al., 2014; Xu et al., 2020), MR diet feeding suppressed the growth of allograft CT26.WT tumors in immunocompromised NSG mice (Figure 2A and 2B, NSG). However, this suppression was lost in immunocompetent Balb/c mice (Figure 2A and 2B, Balb/c). Moreover, although MR diet did not significantly suppress the growth of allograft CT26.WT tumors in Balb/c mice as intestinal tumors in *Apc^min+/-^* mice (Figure 1A-1C), it did induce a significant decrease of circulating CD8^+^CD3^+^ T cells in Balb/c mice (Figure 2C, S2A, and S2B) as observed in *Apc^min+/-^* mice (Figure 1E), along with decreased IFN*γ*^+^CD8^+^ and TNF*α*^+^CD8^+^ T cells (Figure S2C). These observations, together with previous reports that methionine is important for T cell activation and differentiation (Bian et al., 2020; Roy et al., 2020; Sinclair et al., 2019), strongly suggest that dietary methionine restriction may impair the response of tumors to immunomodulators that block the immune checkpoints PD1/PD-L1 or CTLA-4 in T cells (Havel et al., 2019; Robert, 2020).

**Figure 2.**
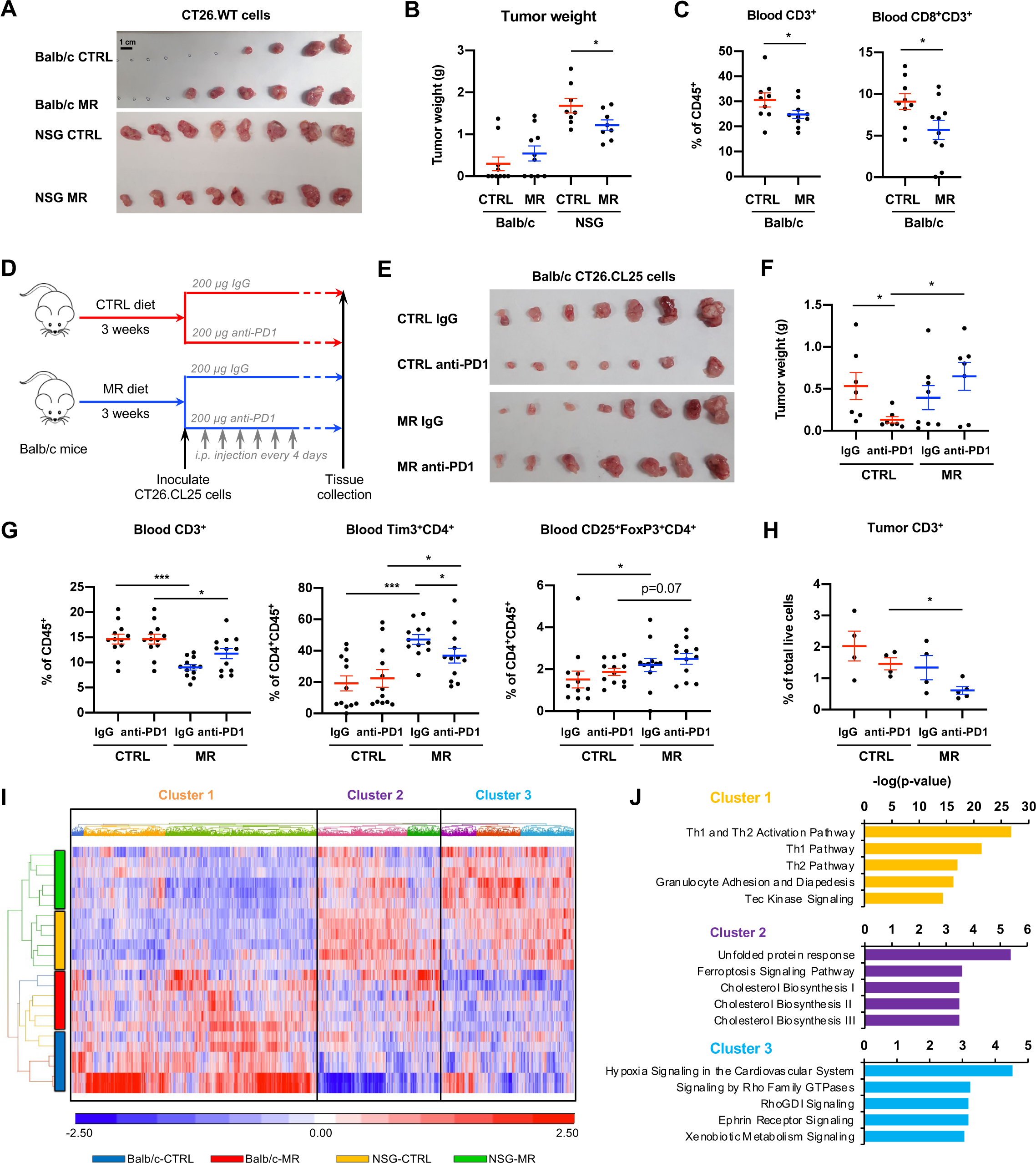
Dietary methionine restriction represses immune activation and blunts tumor response to anti-cancer immunotherapy in immunocompetent mice. (A-B) Dietary methionine restriction represses tumor growth in immunodeficient but not in immunocompetent mice. Immunocompetent Balb/c mice and immunodeficient NSG mice fed with CTRL diet or MR diet were s.c. injected with CT26.WT cells as described in Methods. A final image of the dissected tumors (A) and tumor weights (B) are shown (values are expressed as mean ± s.e.m., n=10 tumors in each group for Balb/c mice; 8 tumors in each group for NSG mice, *p<0.05). (C) Dietary methionine restriction reduces blood CD3^+^ T cells in immunocompetent Balb/c mice. The fractions of blood CD3^+^ and CD8^+^CD3^+^ T cells were analyzed by flow cytometry (values are expressed as mean ± s.e.m., n=10 mice in each group, *p<0.05). (D) Schematic of the anti-PD1 immunotherapy experiment. Immunocompetent Balb/c mice were fed with either CTRL diet or a MR diet for 3 weeks. After s.c. injected with 2×10^5^ CT26.CL25 cells, mice in each group were then randomly divided with 2 groups to be i.p. injected with 200 μg control IgG or anti-PD1 antibody every 4 days. Allografted tumors were monitored and analyzed 2-3 weeks after inoculation. (E-F) Anti-PD1 antibody fails to suppress tumor growth in methionine-restricted immunocompetent mice. A final image of the dissected tumors (B) and tumor weights (C) are shown (values are expressed as mean ± s.e.m., n=7 tumors for CTRL diet groups and n=8 tumors for MR diet groups, *p<0.05). (G) Dietary methionine restriction reduces total circulating CD3^+^ T cells but increases circulating Tim3^+^CD4^+^ exhausted T cells and CD25^+^FoxP3^+^ Treg in Balb/c mice (values are expressed as mean ± s.e.m., n=12 mice in each group, ***p<0.001, **p<0.01, *p<0.05). (H) Dietary methionine restriction reduces total tumor CD3^+^ T cells in anti-PD1 antibody treated mice (values are expressed as mean ± s.e.m., n=4 tumors for CTRL IgG group; 4 tumors for CTRL anti-PD1 group; 4 tumors for MR IgG group; and 5 tumors for MR anti-PD1 group, *p<0.05). (I) Dietary methionine restriction represses immune activation in immunocompetent mice at the molecular level. The transcriptomes of allograft CT26.WT tumors collected from mice treated in (A) were analyzed by RNA-seq as described in Methods (n=6 tumors in each group). The mRNA levels of 8,233 significantly differentially expressed genes were clustered and presented by heatmaps (Cluster 1, 4031; Cluster 2, 2041; Cluster 3, 2151). (J) Dietary methionine restriction differentially altered genes involved in unfolded protein response, ferroptosis, and cholesterol synthesis. The three gene clusters in (I) were subjected to IPA and the top five enriched canonical pathways in each cluster were shown.

To test this possibility, we allografted CT26.CL25 mouse colon carcinoma cells into immunocompetent Balb/c mice under the control diet or MR diet, then injected the mice intraperitoneally with an anti-PD1 antibody or a control IgG (Figure 2D). CT26.CL25 is a subclone of CT26.WT expressing the model tumor-associated antigen beta-galactosidase, which has been widely used as a model for testing immunotherapy protocols and the host immune response (Chen et al., 1996; Irvine et al., 1996; Wang et al., 1995). In line with the observation with CT26.WT cells (Figure 2A), dietary methionine restriction did not significantly affect the growth of allograft CT26.CL25 tumors when treated with control IgG (Figure 2E and 2F, IgG). However, treatment with the anti-PD1 antibody significantly suppressed the tumor growth in control diet-fed Balb/c mice, but it failed to do so in MR diet-fed animals (Figure 2E and 2F, anti-PD1), indicating that dietary methionine restriction indeed reduces the efficacy of immune checkpoint inhibitors (ICI). Further flow cytometry analysis showed that dietary methionine restriction reduced circulating CD3^+^ T cells while increasing Tim3^+^CD4^+^ exhausted T cells and immune suppressive CD25^+^FoxP3^+^ Treg cells (Figure 2G, S2D, S2E, and S2F). Dietary methionine restriction also reduced intratumoral CD3^+^ T cells after the anti-PD1 antibody treatment (Figure 2H and S2G). Therefore, our findings demonstrate that dietary methionine restriction represses T cell activation and dampens the efficacy of ICI-mediated anti-tumor therapy, and further suggest that a sufficient dietary methionine intake is critical to maintain the sensitivity of tumors to anti-tumor immunotherapy in immunocompetent mice.

In support of the notion that the impact of dietary methionine restriction on the anti-tumor immunity is dependent on the host immune system, transcriptomic analysis of allograft CT26.WT tumors collected from Balb/c mice and NSG mice fed with the control diet or MR diet by RNA-seq revealed that the overall transcriptomic profiles of allograft CT26.WT tumors from Balb/c mice were distinct from those of allograft CT26.WT tumors from NSG mice (Table S1, Figure S3A, Balb/c groups vs NSG groups). Gene Set Enrichment Analysis (GSEA) of these transcriptomic datasets further showed that methionine restriction suppressed expression of genes associated with immune system and transportation of small molecules but increased expression of genes associated with cholesterol biosynthesis and TCA cycle in tumors from Balb/c mice (Figure S3B, left). However, the same dietary regimen enhanced immune system (primarily myeloid cells) and transportation of small molecules but suppressed lipid metabolism in tumors from NSG mice (Figure S3B, right), highlighting the importance of these pathways in mediating differential responses of allograft tumors to dietary methionine manipulation in these two mouse strains. Further hierarchical clustering analyses of the total 5,490 differentially expressed genes indicated that tumors from Balb/c mice had higher expression of genes involved in immune functions, including both Th1 and Th2 activation pathways (Figure 2I and 2J, Cluster 1), whereas tumors from NSG mice had higher expression of genes involved in hypoxia and membrane traffic signaling (Figure 2I and 2J, Cluster 3). Genes involved in unfolded protein response, ferroptosis, and cholesterol synthesis were low in allograft tumors from Balb/c mice compared to those from NSG mice (Figure 2I and 2J, Cluster 2). Intriguingly, these pathways were differentially regulated in tumors by dietary methionine in Balb/c mice and NSG mice, as dietary methionine restriction induced their expression in tumors from Balb/c mice but repressed it in tumors from NSG mice (Figure 2I and 2J, Cluster 2). Altogether, these transcriptomic alterations were consistent with our observations that dietary methionine restriction differentially impacted the growth of allograft CT26.WT tumors in Balb/c mice and NSG mice (Figure 2A and 2B), and suggest that host immune activity is a key determinant of the tumor response to dietary methionine intervention and anti-tumor immunotherapy.

### Dietary methionine restriction promotes intestinal cancer progression and suppresses anti-tumor immunity through gut microbiota

Methionine has been reported to promote T cell activation and differentiation through a SAM-mediated cell-autonomous enhancement of histone activation marks, such as H3K4me3 on the promoters of genes involved in cytokine production and cell cycle progression (Roy et al., 2020; Sinclair et al., 2019) or H3K79me2 on key immuno-regulatory transcription factor STAT5 (Bian et al., 2020). However, gut microbiota also plays a fundamental role in the induction, training and function of the host immune system (Belkaid and Hand, 2014) as well as in the efficacy of anti-tumor immunotherapies (Routy et al., 2018; Sivan et al., 2015; Skelly et al., 2019; Vetizou et al., 2015). We hypothesized that in addition to its direct effects on the immune cells, dietary methionine may affect basal systemic immune function, anti-tumor immunity, and outcomes of anti-tumor immunotherapy through modulating gut microbiota.

To test this hypothesis, we first analyzed whether dietary methionine restriction induces any changes of gut bacterial composition in mice. Fecal bacterial 16S rRNA gene amplicon sequencing analysis revealed that fecal bacteria from the MR diet-fed *Apc^min+/-^* mice had a slightly reduced diversity compared to those from the control diet-fed *Apc^min+/-^* mice (Figure S4A). At the phylum level, dietary methionine restriction increased fecal abundance of Firmicutes but decreased Bacteroidetes, resulting in a significant increase of the Firmicutes/Bacteroidetes (F/B) ratio (Figure 3A). Notably, Verrucomicrobia, a phylum of bacteria that in the gut commensal microbiota primarily consists of *Akkermansia muciniphila*, represented an average of 6% of total bacteria in the feces of *Apc^min+/-^* mice fed with the control diet. However, methionine restriction depleted this phylum in almost all mice (Figure 3A, 3B, and S4B). These observations are indicative of a broad impact of dietary methionine on gut microbiota. Dietary methionine restriction also significantly reduced the fecal abundance of *Odoribacter* but increased *Bifidobacterium* in *Apc^min+/-^* mice (Figure 3B, S4B, Table S3). *A. muciniphila*, *Odoribacter*, and *Bifidobacterium* are all major types of gut commensal bacteria frequently associated, negatively and/or positively, with colon carcinogenesis and therapeutic outcomes (Longhi et al., 2020; Saus et al., 2019). In particular, *A. muciniphila* is an abundant mucin-degrading intestinal bacterium (Cani and de Vos, 2017; Derrien et al., 2008) that has a number of beneficial impacts on metabolism and immunity, including improvement of diet-induced obesity in both rodents and humans (Depommier et al., 2019; Everard et al., 2013; Greer et al., 2016) and increase of efficacy of anti-PD1 immunotherapy (Routy et al., 2018). In the gut, the abundance of *A. muciniphila* is reduced in colitis patients (Earley et al., 2019), and *A. muciniphila* supplementation has been shown to repress intestinal inflammation and colitis (Bian et al., 2019; Derrien et al., 2008; Zhai et al., 2019), enhance mucosal wound healing (Alam et al., 2016), and blunt colitis-associated tumorigenesis (Wang et al., 2020). However, despite these studies, how environmental signals modulate the abundance of *A. muciniphila* remains poorly understood. Our observation that *A. muciniphila* is reduced by methionine restriction (Figure 3B) suggests that dietary methionine may be an important nutritional regulator of this bacterium.

**Figure 3.**
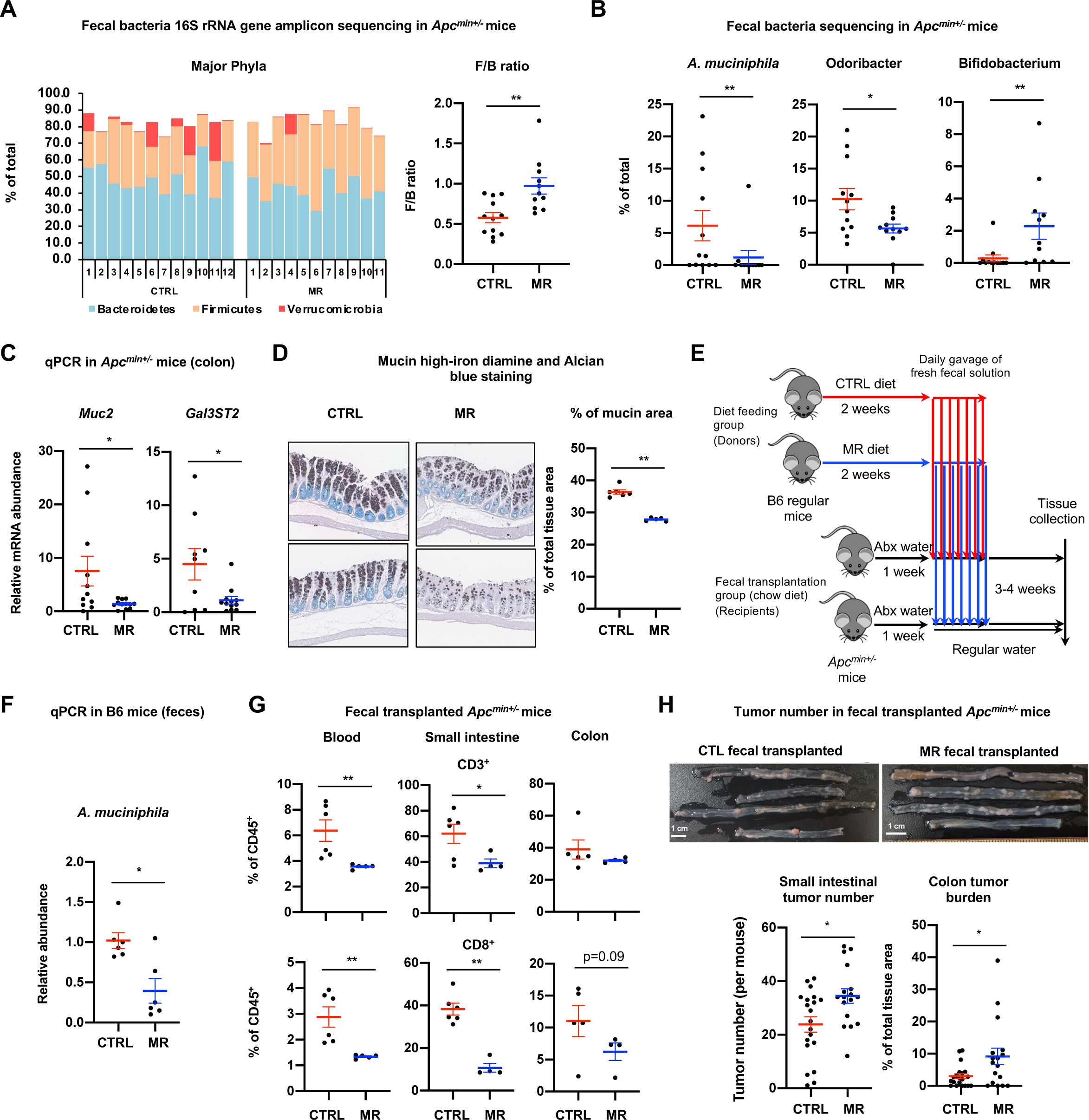
Dietary methionine restriction promotes intestinal cancer progression and suppresses anti-tumor immunity through gut microbiota in immunocompetent mice. (A) Dietary methionine restriction alters gut bacterial composition in *Apc^min+/-^* mice. *Apc^min+/-^* mice were fed with a CTRL diet or a MR diet. Stool DNA samples were analyzed for microbiome using 16S rRNA gene amplicon sequencing as described in Methods (values are expressed as mean ± s.e.m., n=12 mice on CTRL diet and 11 mice on MR diet, **p<0.01). (B) Dietary methionine restriction significantly disrupts three major bacterial groups in *Apc^min+/-^* mice. The fecal abundance of *A. muciniphila, Odoribacter*, and *Bifidobacterium* in diet fed *Apc^min+/-^* mice were determined by 16S rRNA gene amplicon sequencing as in (A) (values are expressed as mean ± s.e.m., n=12 mice on CTRL diet; 11 mice on MR diet, **p<0.01, *p<0.05). (C) Dietary methionine restriction represses the expression of *Muc2* and *Gal3ST2* in *Apc^min+/-^* mice. The mRNA abundance of *Muc2* and *Gal3ST2* was analyzed by qPCR with Lamin as a loading control (values are expressed as mean ± s.e.m., n=6 mice on CTRL diet and 5 mice on MR diet, *p<0.05). (D) Dietary methionine restriction reduces the mucin in the colon of in *Apc^min+/-^* mice. The colonic swiss sections of *Apc^min+/-^* mice on different diets were stained with the high-iron diamine and Alcian blue as described in Methods for mucin protein expression. The % of total mucin positive area (among total tissue area) were quantified in Fiji (values are expressed as mean ± s.e.m., n=6 mice on CTRL diet and 5 mice on MR diet, **p<0.01) (E) Schematic of the fecal transplantation experiment. C57BL/6J mice were fed with either CTRL diet or a MR diet for 2 weeks. Fresh fecal solutions from these fed mice were then gavaged into antibiotic treated *Apc^min+/-^* mice daily for one week. The recipient *Apc^min+/-^* mice were fed with regular chow diet for the duration of the experiment, and intestinal tumors were analyzed 2-3 weeks after fecal transplantation. (F) Dietary methionine restriction reduces *A. muciniphila* in C57BL/6J mice. The relative abundance of *A. muciniphila* in feces of diet-fed C57BL/6J mice (donors) were analyzed by qPCR with total 16S rRNA gene as loading control (values are expressed as mean ± s.e.m., n=6 mice in each group, *p<0.05). (G) *Apc^min+/-^* mice transplanted with feces from MR diet fed C57BL/6J mice have reduced abundance of CD3^+^ and CD8^+^ T cells in the blood and small intestine (values are expressed as mean ± s.e.m., n=6 mice transplanted with feces from mice fed CTRL diet and 4-5 mice transplanted with feces from mice fed MR diet, **p<0.01, *p<0.05). (H) *Apc^min+/-^* mice transplanted with feces from MR diet-fed C57BL/6J mice have more intestinal tumors (values are expressed as mean ± s.e.m., n=20 mice transplanted with feces from mice fed CTRL diet and 17 mice transplanted with feces from mice fed MR diet, *p<0.05).

To better understand how dietary methionine may affect the abundance of *A. muciniphila*, a mucin-degrading bacterium, we analyzed the expression of *Muc2*, a prominent member of mucin family secreted from goblet cells in the colonic epithelium to form the mucus gel layer (Pelaseyed et al., 2014), and *Gal3ST2*, a major galactose-3-O-sulfotransferase responsible for synthesis of colonic sulphomucin (Seko et al., 2002), which is a key substrate of *A. muciniphila* (Derrien et al., 2004; Earley et al., 2019). The mRNA levels of both *Muc2* and *Gal3ST2* were significantly reduced in the colon tissue of methionine-restricted *Apc^min+/-^* mice (Figure 3C). Further staining of colonic mucin also revealed a significant reduction of mucin-positive epithelium in methionine-restricted *Apc^min+/-^* mice (Figure 3D), suggesting that reduced production of colonic mucin, particularly sulphomucin, might be responsible for the limited growth of *A. muciniphila* in these mice.

We next sought to assess the importance of the gut microbiota in mediating dietary methionine-induced modulation of T cell activation and tumor progression in the intestine by microbial repopulation/transplantation experiments. Since *A. muciniphila*-induced T cell responses are limited to T follicular helper cells in a gnotobiotic setting and its impact on intestinal adaptive immune responses is context dependent in conventional mice (Ansaldo et al., 2019), we decided to transplant the whole fecal microbial contents from tumor-free C57BL/6J (B6) mice on either MR diet or control diet to antibiotics-treated *Apc^min+/-^* mice fed on the chow diet (Figure 3E and S4C). We confirmed that the abundance of fecal *A. muciniphila* was significantly reduced in methionine-restricted B6 donor mice (Figure 3F) and this trend was maintained at the early phase after fecal transplantation in the recipient *Apc^min+/-^* mice (Figure S4D). Remarkably, one-week daily transplantation of fecal solution from MR diet-fed tumor-free B6 mice was sufficient to induce a dramatic reduction of CD3^+^ and CD8^+^ T cells in the blood and small intestine in recipient *Apc^min+/-^* mice compared with fecal transplant from B6 mice on a control diet (Figure 3G and S4E). Moreover, these immune alterations were associated with an increased tumor number in the small intestine and tumor burden in the colon (Figure 3H). These observations support our hypothesis that the gut microbiota mediates, at least in part, the influence of dietary methionine on systemic immune function, anti-tumor immunity, and tumor progression.

To further assess the role of the gut microbiota in mediating the impact of dietary methionine on gut anti-tumor immunity, we analyzed total CD45^+^ immune cells in the small intestine of diet-fed tumor-free B6 donor mice (B6-CTRL and B6-MR) and chow-fed fecal recipient *Apc^min+/-^* mice (Apc-FT-CTRL and Apc-FT-MR) by single-cell RNA-seq (scRNA-seq) (Table S4). Based on their individual transcriptomes, CD45^+^ immune cells in the small intestine were clustered into a total of 26 groups (Table S5A and Figure S5A), which could be further categorized into 7 functional groups based on the expression patterns of known marker genes (Figure 4A, S5A, and S5B). In line with our FACS analysis results in MR-fed *Apc^min+/-^* mice (Figure 1F and 3G), MR diet feeding led to reduction of total T cells, particular CD8^+^ effector T cells and central memory T cells (CD8 CMT) in B6 donor mice (Figure 4B and Table S5A, B6). Unexpectedly, total B cell abundance was increased by MR diet feeding. Notably, the MR diet-induced decrease of CD8^+^ effector T cells and CMT cells but increase of B cells in B6 donor mice were transferred to *Apc^min+/-^* mice by fecal transplantation (Figure 4B and Table S5A, Apc-FT), despite the fact that recipient mice were fed regular chow diet. These results indicate that gut microbiota plays a major role in mediating the impacts of dietary methionine on gut immunity. Further analysis of significantly altered genes in different immune cell groups revealed that CD8^+^ effector T cells in both B6-MR and Apc-FT-MR mice had significant downregulation of genes in pathways mediating immune cell differentiation and activation compared to those in B6-CTRL and Apc-FT-CTRL mice, respectively (Figure 4C and Table S6). Moreover, different immune cells from B6-MR and Apc-FT-MR mice also had significantly reduced expression of several genes involved in cytotoxic activity and T cell-mediated immune responses, such as *Gzmb*, *Il2rb, Gzma, Ccl5, or Ccr9* compared to those from B6-CTRL and Apc-FT-CTRL mice (Figure 4D, 4E, and Table S6). Particularly, both of their NKT cells had significantly reduced expression of *Gzma* and both of their CD4^+^ T cells had significantly reduced expression of *Gzmb* (Figure 4E and Table S6). These results indicate a reduced activity of T cells and NKT cells from these mice.

**Figure 4.**
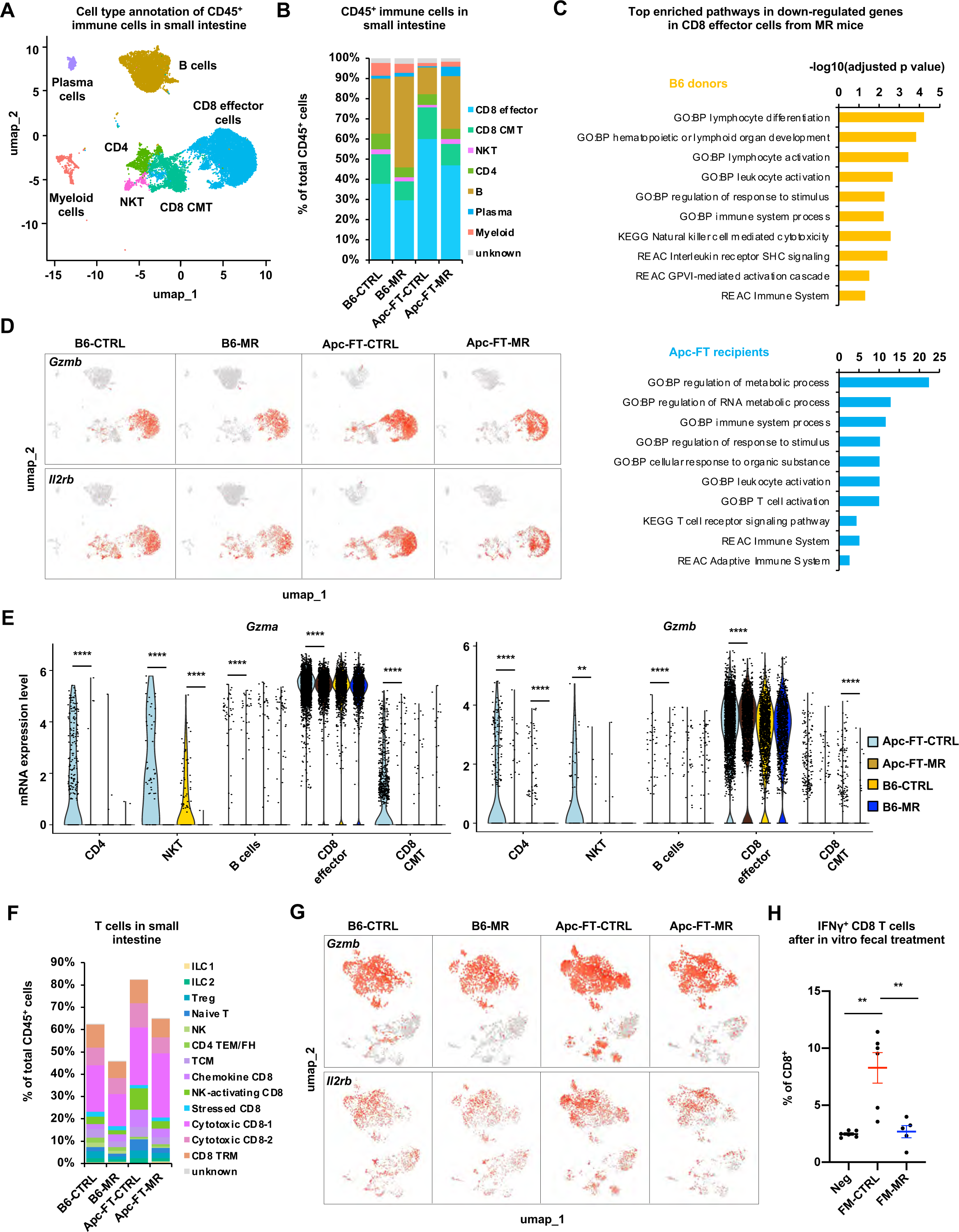
Dietary methionine restriction and MR diet-trained fecal microbiota reduce activation of intestinal T cells. (A) Cell population annotation of CD45^+^ immune cells from CTRL or MR fed B6 donor mice (B6-CTRL or MR) and from *Apc^min+/-^* mice transplanted with fecal solution from CTRL or MR B6 mice (Apc-FT-CTRL or MR). Total CD45^+^ immune cells from the whole small intestine of indicated mice were analyzed by scRNA-seq as described in Methods. (B) Dietary methionine restriction and MR diet-trained fecal microbiota reduces the fraction of CD8^+^ T cells while increasing total B cell population in the small intestine. (C) CD8^+^ effector T cells from MR diet-fed B6 mice and *Apc^min+/-^* mice transplanted with fecal solution from MR B6 mice have reduced expression of genes involved in immune cell function and activation in the small intestine. Significantly downregulated genes in CD8 effector cells in B6-MR mice and Apc-FT-MR mice in Table S6 were analyzed for enriched pathways as described in Methods. Top pathways identified in GO, KEGG, and REAC are shown. (D) Dietary methionine restriction and MR diet-trained fecal microbiota reduces the fraction of expression of *Gzmb* and *Il2rb* in intestinal T cells. Feature plots showing the reduced fraction of *Gzmb^+^* and *Il2rb^+^* T cells in B6-MR and Apc-FT-MR mice. (E) Violin plots showing the reduced expression of *Gzma* and *Gzmb* in different groups of immune cells in B6-MR and Apc-FT-MR mice (**p<0.01, ****p<0.0001). (F) Dietary methionine restriction and MR diet-trained fecal microbiota reduces the fraction of several subgroups of intestinal CD8^+^ T cells. (G) Feature plots showing the reduced fraction of *Ifngr1^+^* and *Il2rb^+^* sub-T cell groups in B6-MR and Apc-FT-MR mice. (H) Fecal microbiota from MR diet fed mice fail to induce the expression of IFN*γ* in CD8^+^ T cells. Purified mouse PBMCs were treated with fecal microbiota (FM) from CTRL diet or MR diet fed donor mice for 12 hours in vitro as described in Methods. The fraction of IFN*γ*^+^ CD8^+^ T cells were analyzed by flow cytometry (values are expressed as mean ±s.e.m., n=5 biological replicates/sample, **p<0.01, student’s t-test).

Additional sub-categorization of T cells based on the expression patterns of known marker genes (Figure S5C and S5D) showed that B6-MR and Apc-FT-MR mice had reduced abundance of central memory T cells (TCM), NK-activating CD8^+^ T cells, tissue resident memory CD8^+^ T cells (CD8 TRM), and certain types of cytotoxic CD8^+^ T cells in their small intestines (Figure 4F, S5E, and Table S5B). The expression of *Gzmb* and *Il2rb* was also reduced in many of these subgroups of T cells (Figure 4G). Finally, when tested for their abilities to activate CD8^+^ T cells *in vitro*, fecal microbes from control diet-fed B6 mice were able to markedly stimulate the expression of IFN*γ* in CD8^+^ T cells after a 12-hour incubation with isolated mouse peripheral blood mononuclear cells (PBMCs), yet fecal microbes from MR diet-fed B6 mice completely failed to do so (Figure 4H and S5F). Taken together, our findings strongly support the notion that dietary methionine exerts a large portion of its impacts on gut anti-tumor immunity through the gut microbiota.

### Dietary methionine restriction reduces fecal hydrogen sulfide production and suppresses anti-tumor immunity

We next investigated the mechanism through which the gut microbiota from methionine-restricted tumor-free B6 mice may suppress anti-tumor immunity and promote tumor growth in recipient *Apc^min+/-^* mice. Although the gut microbiota interacts with the host immune system through multiple complex mechanisms (Belkaid and Hand, 2014), we decided to focus on microbial metabolites, as they are the direct products of dietary interventions. Metabolomics analysis of fecal metabolites showed that dietary methionine restriction significantly altered the abundance of 24 metabolites in B6 donor mice, with 19 of them upregulated (cut off: |fold change|>1.5, p<0.05, Table S6 and Figure S6A). The elevated metabolites were enriched for metabolite sets associated with various types of fatty acid oxidation (Figure S6B), suggesting that dietary methionine restriction alters fatty acid metabolism in the gut. However, very few changes of these metabolites were sustained after 3 weeks of transplantation into the recipient mice (not shown). On the other hand, as expected from the restriction of a sulfur-containing amino acid, fecal H_2_S production activity was significantly blunted in feces from MR diet-fed B6 mice (Figure 5A, B6), and this reduction in H_2_S production activity was preserved in feces from MR fecal recipient *Apc^min+/-^* mice after 3 weeks of transplantation (Figure 5A, *Apc^min+/-^* mice). H_2_S is a gasotransmitter with a wide range of implications in cancer, aging, and age-associated diseases (Cao et al., 2019; Perridon et al., 2016). It is also an endogenous potentiator of T cell activation (Miller et al., 2012). Interestingly, gut is the major H_2_S-producing organ, where the gut microbiota is responsible for about half of this production (Carbonero et al., 2012). Therefore, our observations suggest that dietary methionine might regulate T cell activation through shaping the intestinal microenvironment to support the H_2_S-producing activity of gut bacteria.

**Figure 5.**
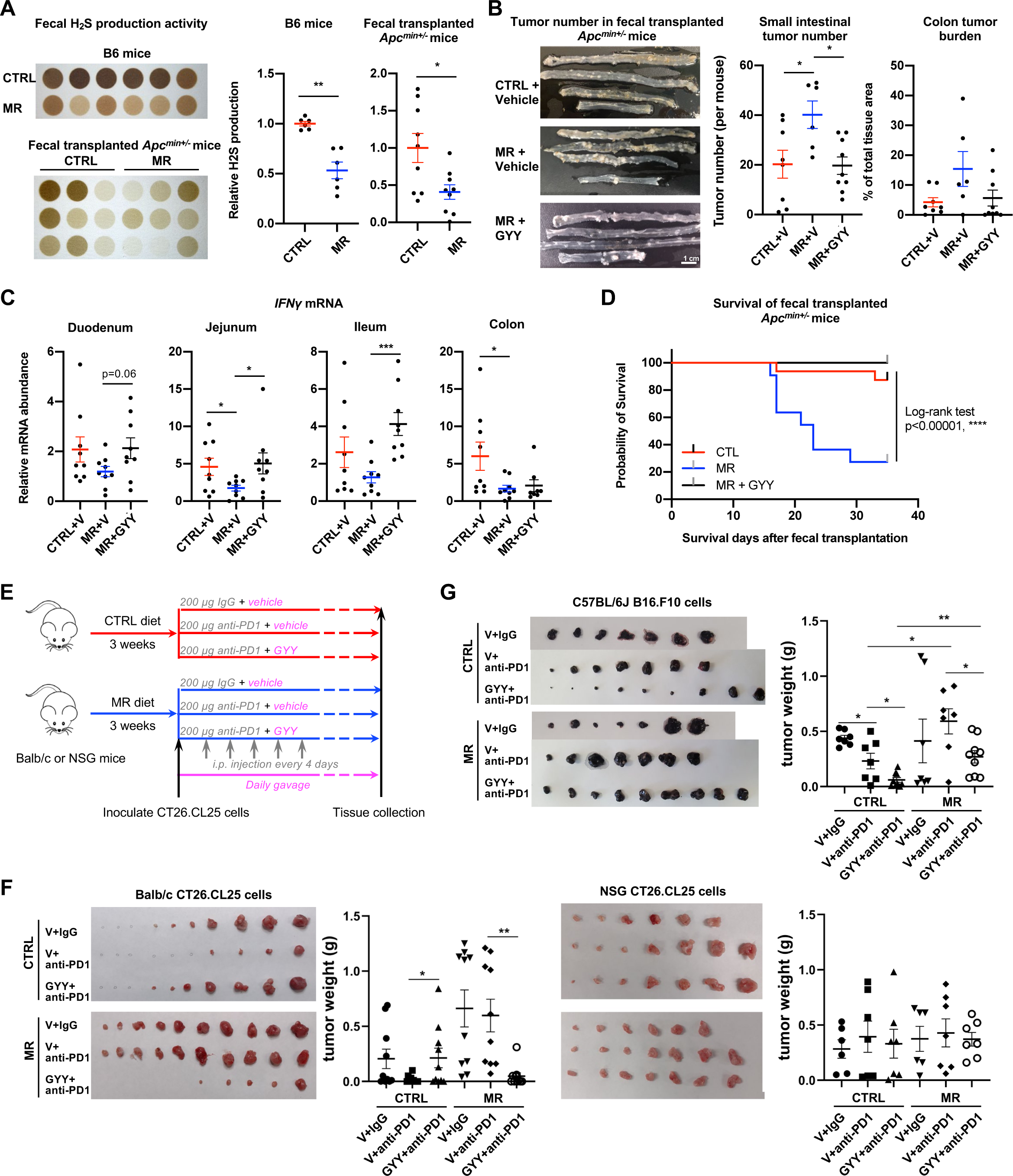
Dietary methionine restriction reduces fecal hydrogen sulfide production and suppresses anti-tumor immunity in immunocompetent mice. (A) Dietary methionine restriction reduces fecal H_2_S production activity. Feces from CTRL or MR diet fed C57BL/6J donor mice as well as respective *Apc^min+/-^* mice recipient mice were subjected to H_2_S production assay using the lead sulfide assay as described in Methods (values are expressed as mean ± s.e.m., n=6 mice each for C57BL/6J donor groups; 9 mice each for *Apc^min+/-^* recipient groups, **p<0.01, *p<0.05). (B) Oral supplementation of GYY4137 rescues MR-induced increase of tumor progression in *Apc^min+/-^* mice. *Apc^min+/-^* mice were fed with CTRL diet or MR diet, together with daily oral gavage of vehicle (V) or GYY4137 (GYY) as described in Methods (values are expressed as mean ± s.e.m., n=8 mice on CTRL diet, 6 mice on MR diet, and 9 mice on MR diet + GYY, *p<0.05). (C) Oral supplementation of GYY4137 rescues MR-induced reduction of *IFNγ* mRNA in the small intestine. *Apc^min+/-^* mice were treated as in (B) and the mRNA levels of *IFNγ* were analyzed by qPCR (values are expressed as mean ± s.e.m., n=9 mice in each group, ***p<0.001, *p<0.05). (D) Oral supplementation of GYY4137 rescues the survival of *Apc^min+/-^* mice on MR diet *Apc^min+/-^* mice were treated as in (B). The health survival of experimental mice was analyzed as described in Methods (values are expressed as mean ± s.e.m., n=16 mice on CTRL diet, 11 mice on MR diet, and 9 mice on MR diet + GYY, ****p<0.0001, Log-rank test). (E) Schematic of the GYY4137 and anti-PD1 immunotherapy experiment. Balb/c or NSG mice were fed with either CTRL diet or a MR diet for 3 weeks. After s.c. injected with 2×10^5^ CT26.CL25 cells, mice in each group were then randomly divided with 3 groups to be daily gavaged with GYY4137, and i.p. injected with 200 μg control IgG or anti-PD1 antibody every 4 days. Allografted tumors were monitored and analyzed 2-3 weeks after inoculation. (F) Oral supplementation of GYY4137 rescues MR-induced resistance of allografted CT26.CL25 tumors to anti-PD1 treatment in immunocompetent mice but not in immunodeficient mice. Balb/c and NSG mice were treated as described in (E). Allografted tumors were monitored and analyzed 2-3 weeks after inoculation (values are expressed as mean ± s.e.m., in Balb/c mice, n=10 tumors for each group; in NSG mice, n=6 tumors for CTRL V+IgG group; 7 tumors for CTRL V+anti-PD1 group; 7 tumors for CTRL GYY+anti-PD1 group; 6 tumors for MR V+IgG group; 7 tumors for MR V+anti-PD1 group, and 7 tumors for CTRL GYY+anti-PD1 group, **p<0.01, *p<0.05). (G) Oral supplementation of GYY rescues MR-induced insensitivity of allografted B16.F10 tumors to anti-PD1 treatment in C57BL/6J mice. Immunocompetent C57BL/6J mice fed with indicated diet and GYY combinations, and then s.c. injected with B16.F10 melanoma cells and i.p. injected with 200 μg control IgG or anti-PD1 antibody as described in Methods (values are expressed as mean ± s.e.m., n=7 tumors for CTRL IgG group; 7 tumors for CTRL anti-PD1 group; 9 tumors for CTRL+GYY anti-PD1 group; 7 tumors for MR IgG group; 7 tumors for MR anti-PD1 group, and 9 tumors for CTRL+GYY anti-PD1 group, **p<0.01, *p<0.05).

To test this possibility, we first examined whether supplementation of GYY4137, a water-soluble slow release H_2_S donor (Lu et al., 2014), could rescue the increase of tumor growth in *Apc^min+/-^* mice transplanted with fecal microbes from MR diet-fed B6 donor mice. As shown in Figure 5B, *Apc^min+/-^* mice transplanted with feces from MR-fed B6 mice had significantly increased tumor number in their small intestine compared to *Apc^min+/-^* mice transplanted with CTRL-fed feces. However, daily oral gavage of GYY4137 significantly reduced the small intestinal tumor number in these mice (Figure 5B, MR+GYY), along with the rescue of the expression of *IFNγ* in the small intestine (Figure 5C) and a striking rescue of their symptom-free survival (Figure 5D), strongly suggesting that defective fecal H_2_S production is responsible, at least partially, for MR diet-induced tumor progression in *Apc^min+/-^* mice.

We next investigated whether supplementation of GYY4137 would rescue MR diet-induced resistance to anti-PD1 immune therapy in allograft CT26.CL25 tumors in immunocompetent Balb/c mice and immunocompromised NSG mice (Figure 5E). Consistent with our observations in Figure 2, allograft CT26.CL25 tumors were sensitive to the treatment of anti-PD1 antibody in Balb/c mice fed with the control diet but resistant to the same therapy when Balb/c mice were fed with the MR diet (Figure 5F, left two panels, V+IgG vs V+anti-PD1). Oral gavage of GYY4137 sensitized allograft CT26.CL25 tumors in Balb/c mice fed with the MR diet to the anti-PD1 treatment (Figure 5F, left two panels, GYY+anti-PD1 in MR). This effect was completely absent in allograft CT26.CL25 tumors on immunodeficient NSG mice (Figure 5F, right two panels), indicating that dietary methionine and GYY modulates tumor growth through anti-tumor immunity. Similar results were observed in B16.F10 melanoma tumors grafted on C57BL/6J mice (Figure 5G), further confirming that MR diet-induced reduction of fecal H_2_S production is responsible for their defective anti-tumor immunity in different mouse strains with independent cancer cell lines.

### Dietary methionine supplementation enhances anti-tumor immunity and suppresses tumor growth in immunocompetent mice

To further confirm the importance of dietary methionine in boosting anti-tumor immunity, we analyzed immunocompetent mice fed with either a chow diet containing 0.4% L-methionine or a custom-made methionine-supplemented chow diet containing 1.3% L-methionine. As expected from our findings with control and MR diets, in C57BL/6J mice, 1.3% methionine diet increased circulating CD3^+^ and CD8^+^ T cells (Figure 6A and 6B). Although the CD4^+^ T cell abundance was not significantly altered (Figure S7A), the population of exhausted CD4^+^ T cells expressing PD-1 and Tim3 was significantly suppressed by this dietary supplementation (Figure 6C). Dietary methionine supplementation further reduced immunosuppressive Treg population in the blood of C57BL/6J mice (Figure 6D). These results indicate that dietary methionine supplementation can indeed boost immunity in immunocompetent animals.

**Figure 6.**
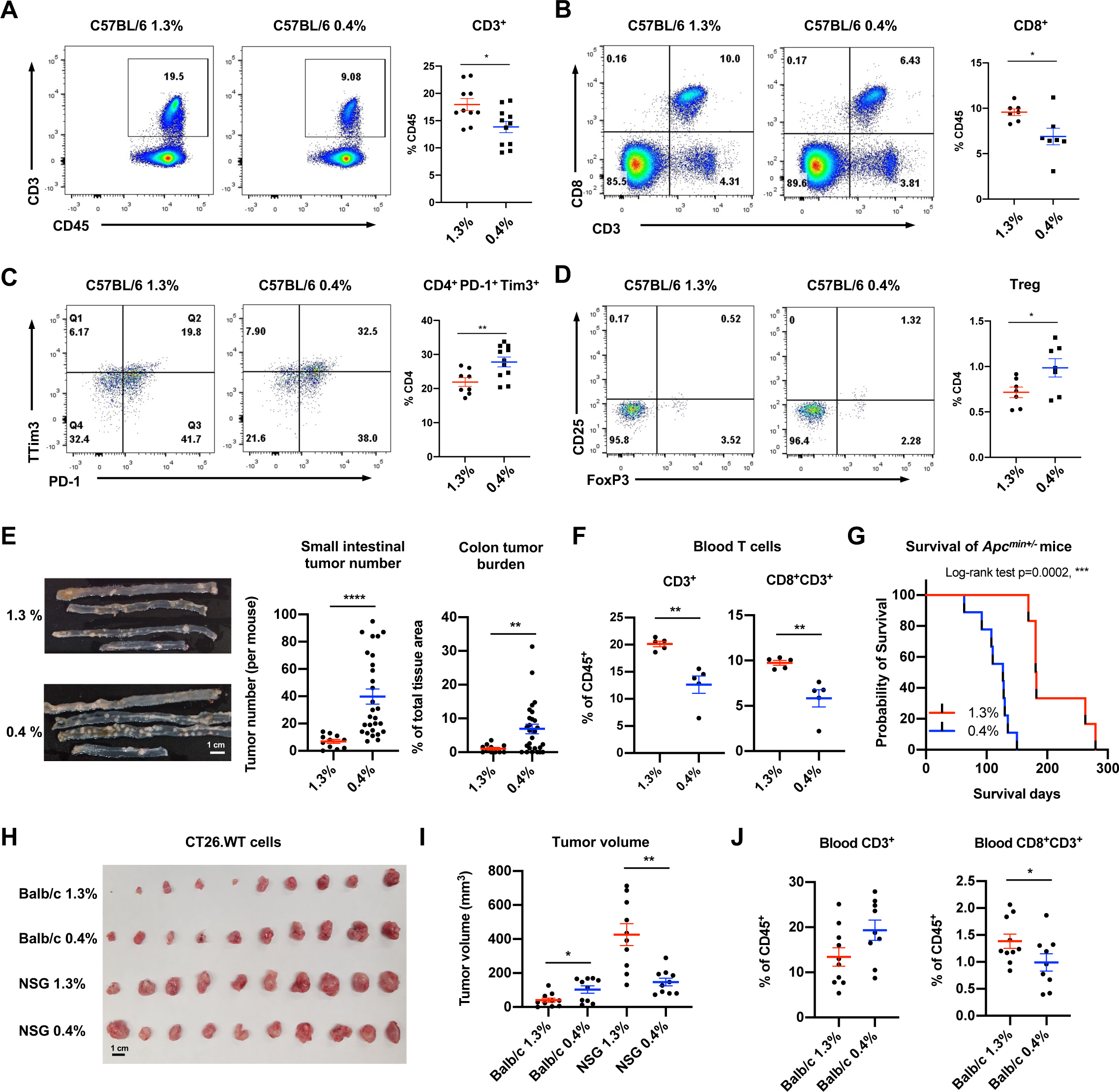
Dietary methionine supplementation activates T cells and suppresses tumor progression in immunocompetent mice. (A) Dietary methionine supplementation increases circulating CD3^+^ T cells in C57BL/6J mice. C57BL/6J mice were fed with chow diet containing 0.4% of methionine and a methionine enriched diet containing 1.3% methionine as described in Methods. The fraction of CD3^+^ T cells in the peripheral blood was analyzed by flow cytometry (values are expressed as mean ± s.e.m., n=10 mice on 1.3% diet and 11 mice on 0.4% diet, *p<0.05). (B) Dietary methionine supplementation increases circulating CD8^+^ T cells in C57BL/6J mice. C57BL/6J mice were fed as in (A). The fraction of blood CD8^+^ T cells was analyzed by flow cytometry (values are expressed as mean ± s.e.m., n=7 mice on each diet, *p<0.05). (C) Dietary methionine supplementation reduces exhaustion of circulating CD4^+^ T cells in C57BL/6J mice. The fractions of CD4^+^PD-1^+^Tim3^+^ T cells in the peripheral blood of mice in (A) were analyzed by flow cytometry (values are expressed as mean ± s.e.m., n=8 mice on 1.3% diet and 11 mice on 0.4% diet, **p<0.01). (D) Dietary methionine supplementation reduces circulating Treg cells in C57BL/6J mice. The fractions of Treg cells in the peripheral blood of mice in (A) were analyzed by flow cytometry (values are expressed as mean ± s.e.m., n=7 mice on each diet, *p<0.05). (E) Dietary methionine supplementation inhibits tumor growth in *Apc^min+/-^* mice. *Apc^min+/-^* mice were fed with chow diet containing 0.4% of methionine and a methionine enriched diet containing 1.3% methionine as described in Methods. Representative gross morphological images of intestinal tumors and total tumor numbers (values are expressed as mean ± s.e.m., n=11 mice on 1.3% diet and 28 mice on 0.4% diet, **p<0.01, ****p<0.0001). (F) Dietary methionine supplementation increases fraction of CD3^+^ and CD3^+^CD8^+^ T cells in the blood of *Apc^min+/-^* mice. The fractions of indicated cell population in the peripheral blood were analyzed by flow cytometry as described in Methods (values are expressed as mean ± s.e.m., n=5 mice in each group, **p<0.01). (G) Dietary methionine supplementation increases survival of *Apc^min+/-^* mice. The survival of *Apc^min+/-^* mice under the indicated diets were monitored (values are expressed as mean ± s.e.m., n= 9 mice on 0.4% diet and 6 mice on 1.3% diet). (H-I) Dietary methionine supplementation represses tumor growth in immunocompetent mice but enhances tumor growth in immunodeficient mice. Immunocompetent Balb/c mice and immunodeficient NSG mice fed with chow diet (0.4%) or a methionine enriched diet (1.3%) were s.c. injected with CT26.WT cells as described in Methods. A final image of the dissected tumors (H) and tumor volumes (I) are shown (values are expressed as mean ± s.e.m., n=10 tumors for each group, *p<0.05, **p<0.01). (J) Dietary methionine supplementation increases blood CD8^+^CD3^+^ T cells in Balb/c mice. The fractions of blood CD3^+^ and CD8^+^CD3^+^ T cells were analyzed by flow cytometry (values are expressed as mean ± s.e.m., n=10 mice on 1.3% diet and 9 mice on 0.4% diet, *p<0.05).

In support of this notion, dietary methionine supplementation reduced the intestinal tumor number and burden in *Apc^min+/-^* mice (Figure 6E), increased the fractions of circulating CD3^+^ and CD8^+^CD3^+^ T cells (Figure 6F and S7B), and dramatically extended their health-span (Figure 6G). Remarkably, compared to 0.4% chow diet, 1.3% methionine-supplemented diet suppressed the growth of allograft CT26.WT tumors in immunocompetent Balb/c mice but markedly enhanced their growth in immunocompromised NSG mice (Figure 6H and 6I). Such suppression of tumor growth in Balb/c mice fed high-methionine diet was coupled with a significant increase of circulating CD8^+^CD3^+^ T cells (Figure 6J and S7C). In sum, our findings indicate that dietary methionine supplementation enhances anti-tumor immunity in immunocompetent individuals but promotes tumor growth in immunocompromised animals.

## Discussion

As a sulfur-containing proteinogenic amino acid, methionine has a broad array of cell autonomous impacts on protein synthesis, histone methylation, and redox homeostasis in both cancer cells and immune cells, thereby is essential for their proliferation, stress resistance, and overall functions (Gao et al., 2019; Mentch et al., 2015; Roy et al., 2020; Sinclair et al., 2019; Wang et al., 2019; Xu et al., 2020). As such, methionine in tumor microenvironment has been shown to directly regulate the interaction between cancer cells and immune cells and influence anti-tumor immunity (Bian et al., 2020). The net effect of dietary methionine on cancer growth, therefore, depends on the relative dependence of cancer cells vs immune cells on this nutrient. In the present study, we demonstrate that in addition to the reported cell-autonomous effects, the impact of dietary methionine on tumor growth and anti-tumor immunity is also dependent, at least in part, on gut microbiota-mediated non-cell autonomous activation of immune cells. We show that while repressing tumor growth in immunodeficient mice, dietary methionine restriction enhances tumor growth in immunocompetent animals. We provide evidence that this tumor-promoting effect of methionine restriction in immunocompetent mice is due to impaired T cell activation, which also dampens the efficiency of ICI-mediated anti-tumor immunotherapy. We further show that dietary methionine restriction-induced gut microbiota alteration and subsequent reduction of fecal H_2_S production partially mediate its influence on systemic immune function, anti-tumor immunity, and tumor progression. Collectively, our findings not only highlight the importance of immune activation in the tumor-suppressing action of methionine, but also identify gut microbiota as a key non-cell autonomous link that couples dietary methionine to T cell activation and anti-tumor immunity.

Our study has several important translational implications. Firstly, the observation that dietary methionine restriction suppresses tumor growth in immunodeficient mice but promotes tumor growth and dampens therapeutic response to anti-tumor immunotherapy in immunocompetent mice suggests that in the clinical setting, methionine restriction may be beneficial for treatment of immunocompromised cancer patients but will be detrimental for immunocompetent individuals. Conversely, dietary methionine supplementation may boost anti-tumor immunity in immunocompetent patients but will support tumor growth in immunocompromised cancer patients. Secondly, the discovery that the gut microbiota mediates the influence of methionine restriction on anti-tumor immunity and tumor progression supports the idea that manipulation of the microbiome could be applied for development of new therapeutic anti-cancer approaches. Finally, the positive association between dietary methionine, the H_2_S producing activity of gut microbiota, and anti-tumor immunity revealed in our study raises an interesting possibility that H_2_S chemical donors may help to boost anti-tumor immunity and/or the efficacy of anti-tumor immunotherapies without direct impact on tumor cell proliferation. In support of this notion, GYY4137 has been shown to exhibit anti-tumor activity both *in vitro* and *in vivo* (Lee et al., 2011).

Our study expands our understanding on how dietary methionine regulates the interaction between cancer cells and immune cells indirectly through gut microbiota, which could be employed to development of novel therapeutic strategies against human cancers. However, several questions remain to be answered. First, while we showed that dietary methionine restriction reduces the colonic expression of *Muc2* and *Gal3ST2*, proteins important for formation of colonic mucus gel layer and maintenance of colonic microenvironment for resident gut microbiota such as *A. muciniphila* (Derrien et al., 2004; Earley et al., 2019; Pelaseyed et al., 2014; Seko et al., 2002), future studies are needed to investigate how methionine regulates the expression these enzymes in intestinal epithelium. Second, while we discovered that dietary methionine restriction reduces the H_2_S producing activity of gut microbiota, additional metagenomic and metatranscriptomic analyses are required to address how this dietary manipulation shapes the composition of gut microbiota and/or transcriptionally modulates microbial sulfur metabolism. Last, how H_2_S promotes anti-tumor immunity is still not clear. H_2_S-mediated protein cysteine sulfhydration has been reported to augment GAPDH activity and NF-*κ*B-mediated cell survival (Mustafa et al., 2009; Paul and Snyder, 2012), whereas many other metabolites (e.g. fumarate and itaconate)-induced cysteine modifications on metabolic enzymes and inflammatory proteins suppress immunity (Diskin et al., 2021; Kornberg et al., 2018; Liao et al., 2019; Mills et al., 2018; Qin et al., 2019). Thus, it is possible that H_2_S can help to enhance glycolysis and survival of activated immune cells and/or prevent the accumulation of immune-inhibiting cysteine modifications, which in turn maintain a suitable level of anti-tumor immunity. It will be of great interest to test these possibilities in the future.

In summary, our study reveals unexpected opposing impacts of dietary methionine restriction on tumor progression and therapeutic response in immunodeficient and immunocompetent mice, and further identifies an important role of gut microbiota in mediating the non-cell autonomous influence of dietary methionine on anti-tumor immunity. Our findings may pave the way for new anti-cancer therapeutic strategies, and therefore, have important translational implications in the emergent field of nutritional oncology.

## Methods

### Animal experiments

All mice were maintained under strict SPF conditions at the National Institute of Environmental Health Sciences (NIEHS) animal facility. All mice were housed in micro-isolator static cages (Techniplast, Exton, PA) with autoclaved nesting material (Nestlet, Ancare Corp., Bellmore, NY) and housed on hardwood bedding (Sani-chips, PJ Murphy, Montville, NJ). Methionine restricted diet containing 0.172% DL-Methionine (#510029) and a control diet containing 0.86% DL-Methionine (#510027) were purchased from Dyets, Inc. (Bethlehem, PA). The custom-made 1.3% methionine enriched chow diet and normal chow diet (NIH-31) were purchased from Harlan Teklad. All diets were tested by Quality Analysis Laboratory before feeding.

All animal procedures were reviewed and approved by NIEHS Animal Care and Use Committee. All animals were housed, cared for, and used in compliance with the Guide for the Care and Use of Laboratory Animals and housed and used in an Association for the Assessment and Accreditation of Laboratory Animal Care, International (AAALAC) Program.

### Methionine diet feeding procedure

Six- to eight-week-old *Apc^min+/-^* mice (Jax: 002020) were fed with diets containing different methionine ad libitum for up to 8 weeks with free access to water. The first pair includes a methionine restricted (MR) diet containing 0.172% DL-Methionine or a control diet containing 0.86% DL-Methionine, and the second pair includes the NIH-31 chow diet containing 0.4% methionine and a 1.3% methionine supplemented chow diet. After 8-weeks of feeding, mice were euthanized by CO2 for tissue harvest, and bled immediately afterward for other experiments.

To analyze the long-term impact of methionine diets on symptoms-free survival of *Apc^min+/-^* mice, a small group of *Apc^min+/-^* mice were maintained on their respective diets. Their health status was monitored twice a week, and mice were removed from the study (euthanized) when they lost 20% of their starting body weight, experienced severe rectal bleeding, or displayed any signs of distress such as hunching, labored breathing, incoordination, seizures, lethargy, or becoming prostrate.

### Allograft experiments and PD1 treatment

Six- to eight-week-old BALB/c (Jax: 000651) and NSG (Jax: 005557) mice were fed with two groups of high-methionine and low-methionine diets as described above for 3 weeks. They were then inoculated subcutaneously with 2×10^5^ mouse colon cancer CT26. WT cells and fed with the different methionine diets respectively was continued for an additional 3-4 weeks.

For the immunotherapy experiments with anti-PD1 antibody, the mice were first fed with different methionine diets for 3 weeks, and then inoculated subcutaneously with 2×10^5^ mouse colon cancer CT26.CL25 cells. Isotype-matched control antibody rat IgG2a (BioXCell, BE0089) and rat anti-PD1 antibody (BioXCell, BE0146) were given intraperitoneally at a dose of 200 μg per mouse on day 4 after cell inoculation, then every 4 days for the duration of the experiment. Mice were continually fed with their respective diets for about additional 3-4 weeks with free access to water. No mouse had 20% loss of initial body weight, severe abdominal distension, tumor size reached 1.5 cm^3^ or interferes with normal ambulation or ulcerates.

### Fecal transplantation procedure

One cohort of 6-8 weeks old C57BL/6J mice fed with control or MR diets (6 mice/diet) for 2 weeks were used as donors to provide fresh feces for fecal transplantation. Gut microbiota depletion and fecal transplantation were conducted as follows: 8-10-week-old *Apc^min+/-^* mice fed on regular chow diet were given autoclaved RO/DI water containing an antibiotic cocktail (1 g/L Ampicillin and 0.575 g/L Enrofloxacin) for 1 week, were then inoculated daily by oral gavage for 7 days with 100 μl/mouse of fecal solution prepared with combined feces freshly collected from donor C57BL/6J mice (1:1 dilution of feces with PBS). *Apc^min+/-^* mice were maintained with sterile RO/DI water and chow diet during fecal transplantation and afterward. They were monitored for their health status as described above daily, then analyzed for intestinal tumor formation 3-4 weeks after the fecal transplantation.

### GYY4137 supplementation procedure

Six-eight weeks old C56BL/6J, BALB/c mice, and immunodeficient NSG mice were divided into six groups each, with three groups fed with the control diet and three groups fed with the MR diet for 3 weeks. Starting from Day 21, IgG and one anti-PD1 groups were dosed daily with a vehicle carboxymethyl cellulose (CMC), the other ani-PD1 group were dosed daily with 50 mg/Kg GYY4137, respectively through gavage. Also, on Day 21, they were inoculated (s.c.) with 2×10^5^ cancer cells (the murine melanoma cell line B16.F10 or the murine colon carcinoma cell line CT26.CL25) and administered with 200 μg IgG or anti-PD1 per mouse on day 4 after cell inoculation, then every 4 days for the duration of the experiment. The mice were continually fed with their respective diets for about additional 4 weeks with free access to water. For fecal transplantation experiment, CTRL group and the MR group were gavaged daily with 100ul water, and the other MR group were gavaged daily with 50 mg/Kg GYY4137 during fecal transplantation and afterward.

### Image and Immunohistochemistry quantification

Total tumor numbers in the small intestine of *Apc^min+/-^* mice under different treatments were counted manually. The tumor surface area in the colon and the colonic tissue surface area were quantified in Fiji (ImageJ v.2.1.0/1.53g) and the final tumor burden was calculated using the total surface area of tumors against the total surface area of the colon.

The Ki67 positive area in the ileum and the mucin positive area in the colon of *Apc^min+/-^* mice under control or MR diets were quantified in Fiji using “Colour Deconvolution” with vector of H DAB. The final percentage of positive area was calculated using the total staining area against the total tissue area.

### Immunoblotting and quantitative real-time PCR

Proteins were separated using a 4–20% Mini-PROTEAN® TGX™ Precast Protein Gels and transferred onto Immobilon-FL PVDF membrane (Millipore). The membranes were blocked in 5% BSA for 1 hour, and then incubated with a specific primary antibody at 4 °C overnight. Blots were probed with anti-Bax (Cell Signaling, cs2772, 1:1000) and anti-beta-actin (Millipore Sigma, MAB1501, 1: 10,000) antibodies. The membranes were then washed with 1xTBS-T (0.1%Tween20) with vigorous shaking on a platform shaker at room temperature for 4×5 minutes. The membranes were further probed with secondary antibody IRDye 800CW (1:20,000) for 1 hour. After rinse, the membranes were scanned on an Odyssey imaging system.

Total RNA was purified from tissues using RNeasy Mini kit (Qiagen) followed by cDNA synthesis with the High-Capacity cDNA Reverse Transcription Kit (ThermoFisher Scientific). Real-time PCR was performed using iQ SYBR Green Supermix (Biorad). Specific primers are listed in Table S8.

### Flow cytometry analysis

Blood samples were collected with EDTA coated tubes. Red blood cells were lysed with ACK lysis buffer at room temperature for 10 min. The collected lymphocytes were incubated with anti-mouse CD16/32 at room temperature for 10 min to block the IgG Fc receptors. Expression of surface markers was detected by simultaneously staining with the following antibodies (eFluor 450 anti-mouse CD45, BV510 anti-mouse CD45, APC-Cy7 anti-mouse CD4, FITC anti-mouse CD4, APC anti-mouse CD25, PerCP-eFluor 710 anti-mouse CD3, FITC anti-mouse CD3, PE-Cy7 anti-mouse CD3, FITC anti-mouse CD8, PerCP-Cy™5.5 anti-mouse CD8, PE anti-mouse CD279 (PD-1), BV711 anti-mouse Tim3) and LIVE/DEAD fixable Aqua dead cell stain kit (ThermoFisher, Cat. # L34957) at room temperature for 15 min followed by flow cytometry.

For FoxP3 staining, after perform cell surface staining, the cells were fixed with True-Nuclear™ Transcription Factor Fixation Buffer (Biolegend, Cat. # 424401) at room temperature for 45 min. Add 2 ml PBS to wash cells once. Add 1 ml the True-Nuclear™ 1x Perm Buffer to each tube, centrifuge tubes at 400 g at room temperature for 5 minutes and discard the supernatant. Resuspend the cell pellet in 100 µL of 1x Perm buffer with antibody for 30 minutes in the dark. Add 1 ml of 1x Perm buffer to each tube. Centrifuge the tubes at 400g at room temperature for 5 minutes and discard the supernatant. Add 1 ml cell staining buffer (Biolegend, Cat. # 420201) to each tube. Centrifuge the tubes at 400g at room temperature for 5 minutes and discard the supernatant. Resuspend in 0.3ml cell staining buffer, and then acquire the tubes on a flow cytometer. The IFNγ and TNFα staining were performed using a similar protocol, except that cells were first fixed using BD Cytofix ™ solution (Cat. # 554655) then permeabilized using BD Perm/Wash buffer (Cat. # 554723). Flow cytometric analysis was performed on BD LSRFortessa instrument (BD Biosciences) and analyzed using FACSDiva (BD Biosciences) or FlowJo V10 (FlowJo LLC) software. All antibodies are listed in Table S9.

### Immunohistochemistry

Immunohistochemical staining for Ki-67 was performed using the HRP-polymer technique. Formalin-fixed, paraffin-embedded tissue sections on glass slides were deparaffinized in xylene and rehydrated through graded ethanol. Heat-induced epitope retrieval was performed using a citrate buffer (pH6.0, Biocare Medical, Concord, CA) in the Decloaker® pressure chamber for 15 minutes at 110 °C. Then endogenous peroxidase blocking was done by immersing the sections in 3% H2O2 for 15 minutes. Non-specific sites were blocked by incubating slides for 20 minutes with Rodent Block M (Biocare Medical, Concord, CA). The sections were then incubated with rabbit anti-Ki-67 antibody (Cat # ab16667, Lot # GR3341233-10, Abcam, 0.05 mg/ml) at a 1:100 dilution for 60 minutes at room temperature. No negatives were run. The antigen-antibody complex was detected using Rabbit on Rodent HRP Polymer (Biocare Medical, Concord, CA) and 3,3-diaminobenzidine (Dako, Carpinteria, CA). Slides were then counterstained with hematoxylin hydrated and coverslipped.

### RNA-seq analysis

RNA was extracted from allografted CT26.WT tumors of mice with QIAzol, and purification of total RNA was performed with the QIAGEN RNeasy RNA isolation kit (Redwood City, CA) according to the manufacturer’s protocol. RNA-seq libraries were generated with 1 μg of RNA as input using the TruSeq RNA Sample Prep Kit (Illumina, San Diego, CA) with poly(A)-enrichment according to the TruSeq protocol. Index multiplexed samples were sequenced using the 75-bp single-end protocol via the NextSeq500 (Illumina) according to the manufacturer’s protocol. Reads (30–60 million reads per sample) were aligned to the University of California Santa Cruz (UCSC) mm10 reference genome using STAR (Version 2.6) with Gencode vM23 annotation. The quantification results from “featureCount (subread, Version 1.4.6)” were then analyzed with the Bioconductor package DESeq2, which fits a negative binomial distribution to estimate technical and biological variability. Comparisons were made between experimental conditions and a gene was considered differentially expressed when the p value for differential expression was less than 0.01.

Significant gene lists were subjected to Ingenuity Pathway Analysis (IPA, QIAGEN, USA; Content version 60467501). GSEA was performed using GSEA software (Version 4.1). Genes were pre-ranked based on the fold change of gene expression. This application scores a sorted list of genes with respect to their enrichment of selected functional categories (Kyoto Encyclopedia of Genes and Genomes [KEGG], Biocarta, Reactome, and gene ontology [GO]). The significance of the enrichment score was assessed using 1000 permutations. Benjamini and Hochberg’s false discovery rate (FDR) was calculated for multiple testing adjustments. q-value < 0.05 was considered significant. The resulting enriched pathways were visualized using the Cytoscape (Version 3.8.2) Enrichment Map plugin.

### scRNA-seq analysis of small intestinal CD45^+^ immune cells

#### Isolation of CD45^+^ immune cells from the small intestine

To analyze total CD45^+^ immune cells from diet-fed C57BL6/J donor mice and fecal transplanted *Apc^min+/-^* mice, mice were euthanized and surface-sterilized using 70% (vol/vol) ethanol. After removing the mesentery/fat and Peyer’s patches, the small intestine was flashed with ice cold PBS (Mg^2+^Ca^2+^ free), then opened longitudinally and removed the mucous layer by gently rubbing the intestine between fingers in cold PBS. The small intestine was cut into 0.5-1cm pieces and washed twice in 20 ml PBS (Mg^2+^Ca^2+^ free) containing 0.37 mg/ml EDTA (1.25 mM) and 0.145 mg/ml DTT (1 mM) in a shaking incubator at 37°C for 15 min, and then twice with 5 ml ice cold PBS (Mg^2+^Ca^2+^ free) by gently shaking. The remaining tissue was then digested by incubating in digestion medium (20 ml RPMI-1640 containing 5% fetal calf serum, 20 mM HEPES, 100 U/ml penicillin /streptomycin, 1 mg/ml Collagenase I (Sigma-Aldrich, C0130), 1 mg/ml Dispase II (Sigma, D4693) and 20 μg/ml DNase I (Roche, 11284932001)) for 15 - 20 min at 37°C while gently shaking, the liberated cells in the supernatant were collected and placed on ice. Repeat the digestion step for 2 additional times with 10-15 ml fresh digestion medium each time. Following the third digestion step all fractions were combined and passed through a 70 um strainer (Corning, 431751). The filtered single cells were collected, washed with 10 mL ice cold sterile PBS (without FCS) once, and resuspend in 100 μl ice cold sterile PBS. The resulting single cell suspension was stained for CD45^+^ and PI, and sorted by flow cytometry. The final CD45^+^ single cells were used for generating scRNA-seq libraries.

#### scRNA-seq library preparation and sequencing

The cells were counted and examined for viability with trypan blue staining using a TC-20 cell counter (Bio-Rad). Approximately 10,000 live cells with above 70% viability were loaded into the Single Cell Chip to generate single cell emulsion in Chromium Controller (10x Genomics, 120263) with Chromium Single Cell 3’ Library & Gel Bead Kit v3.1 (10x Genomics, 1000268). Reverse transcription of mRNA and cDNA amplification were carried out following the manufacture’s instruction (10x Genomics, 1000268). The amplified cDNA was further fragmented to construct NGS libraries. The libraries were then sequenced by the NIEHS Epigenomics and DNA Sequencing Core Laboratory with the parameters recommended in the manufacture’s instruction.

#### scRNA-seq data processing

Raw read processing was carried out using the Cell Ranger Single-Cell Software Suite (version 6.0.1, 10x Genomics Inc., CA). Briefly, the demultiplexed FASTQ files (paired-end, Read 1: 30bp, Read 2: 100bp) were generated using the CellRanger *mkfastq* command. The primary data analyses which included alignment, filtering, barcode counting and UMI quantification for determining gene transcript counts per cell (generated a gene-barcode matrix), quality control, clustering and statistical analysis were performed using CellRanger *count* command with genome reference “refdata-gex-mm10-2020-A”.

Raw gene expression matrices generated per sample by CellRanger were imported into R (version 4.0.1) and converted to a Seurat object using the Seurat R package (version 4.0.2) (Butler et al., 2018). Function *decontX* from R package “celda” (Campbell et al., 2021) was used to remove RNA contamination. Dead cells and doublets were also removed. Low quality cells were further filtered out based on the following steps. First, the total number of UMIs and genes, and percentage of UMIs derived from mitochondrial genome for each cell were counted. Then, the upper bound was calculated as mean plus two standard deviation (SD) and the lower bound as mean minus two SD for both the total UMIs and genes, respectively. Next, cells which had over 10% UMIs derived from mitochondrial genome were discarded. Finally, cells with total UMIs or genes outside of the upper and lower bounds were removed.

### Single-cell gene expression quantification and major cell type classification

For the remaining cells (cleaned cells), Seurat *SCTransform* function was used for gene expression normalization, highly variably genes (HVG) and scaling. Gene expression data were normalized to total cellular read count(10000/cell); and the top 3000 highly variably genes (HVGs) were used as features for dimensionality reduction and clustering. Cell-Cycle scores were also calculated using Seurat *CellCycleScoring* function from a set of cell-cycle related gene pairs as input to classify cells into G1, S and G2/M phase. Cell cycle effect was regressed out using *SCTransform* function as well. The Seurat *RunPCA* functions was performed to calculate PCA and top 40 principal components (PCs) selected for downstream analysis. After R package “Harmony” (Korsunsky et al., 2019) corrected the batch effects, the Seurat FindNeighbors function was used to construct a Shared Nearest Neighbor (SNN) Graph, and the RunUMAP function was then applied to plot the selected significant PCs. FindClusters function with “resolution = 1” parameter was carried out to cluster cells into different groups.

To identify marker genes for each cluster or cell type, the Seurat *FindMarkers* function was applied to compare the gene expression values of cells from the cluster of interest to that of cells from the rest of clusters. Cell clusters were annotated by the following two approaches: 1) the canonical marker genes were applied to annotated cell clusters to known biological cell types; 2) the predicted marker genes were applied for each cluster. The predicted marker genes could further confirm the annotated cell type, at the same time, they could also be used to name cell clusters which don’t have canonical marker genes.

Seurat *FindMarkers* function with parameter of “MAST” (Finak et al., 2015) test was used to calculate DEG between two group cells of interest. The final DEG lists in Table S6 were generated using cut offs of padj<0.05 and/or |FC|>1.3. The downregulated pathways enriched in intestinal CD8 effector cells in response to diet feeding or fecal transplantation were analyzed using the corresponding DEG lists in https://biit.cs.ut.ee/gprofiler/gost.

### Analysis of T cell subpopulation

To further explore features of T cells, we carried out a series of analyses focusing on CD8^+^ and CD4^+^ T cells. The T cells were extracted from whole single cell population based on the cell barcodes. Then, gene expression matrix of these T cells was obtained from cleaned data. The downstream analyses, including normalization, HVG selection, scaling, PCA calculation, batch effect correction, clustering and DEG calling were applied following the same procedure as single cell analysis described in previous section.

### Fecal bacterial 16S rRNA gene amplicon sequencing

#### Stool Samples DNA Isolation

Samples were transferred to a 2 ml tube containing 200 mg of ≤106 μm glass beads (Sigma, St. Louis, MO) and 0.5 ml of Qiagen PM1 buffer (Valencia, CA). Bead beating was performed for 5 minutes in a Qiagen TissueLyser II at 30Hz. After centrifugation for 5 minutes, 0.45ml of supernatant was transferred to a new tube containing 0.15ml of Qiagen IRS solution followed by incubation at 4°C overnight. After a brief centrifugation, supernatants were transferred to deep well plates containing 0.45ml of Qiagen binding buffer supplemented with Qiagen ClearMag Beads. DNA was purified using the automated KingFisher™ Flex Purification System and eluted in DNase free water (Imane Allali, 2017; Lucía Guadamuro, 2019; M. Andrea Azcarate-Peril and Schottel, 2018).

#### 16S rRNA gene amplicon sequencing

12.5 ng of total DNA was amplified using universal primers targeting the V4 region of the bacterial 16S rRNA gene. Primer sequences contained overhang adapters appended to the 5’ end of each primer for compatibility with Illumina sequencing platform. The primers used were F515/R806 (J. Gregory Caporaso, 2011). Master mixes contained 12.5 ng of total DNA, 0.5 µM of each primer and 2x KAPA HiFi HotStart ReadyMix (KAPA Biosystems, Wilmington, MA). Each 16S rRNA gene amplicon was purified using the AMPure XP reagent (Beckman Coulter, Indianapolis, IN). In the next step each sample was amplified using a limited cycle PCR program, adding Illumina sequencing adapters and dual-index barcodes (index 1(i7) and index 2(i5)) (Illumina, San Diego, CA) to the amplicon target. The final libraries were again purified using the AMPure XP reagent (Beckman Coulter), quantified and normalized prior to pooling. The DNA library pool was then denatured with NaOH, diluted with hybridization buffer and heat denatured before loading on the MiSeq reagent cartridge (Illumina) and on the MiSeq instrument (Illumina). Automated cluster generation and paired–end sequencing with dual reads were performed according to the manufacturer’s instructions.

#### Bioinformatics Analysis

Sequencing output from the Illumina MiSeq platform were converted to fastq format and demultiplexed using Illumina Bcl2Fastq 2.18.0.12. The resulting paired-end reads were processed using QIIME 2 2018.11 (Bolyen et al., 2019). Index and linker primer sequences were trimmed using the QIIME 2 invocation of cutadapt. The resulting paired-end reads were processed with DADA2 through QIIME 2 including merging paired ends, quality filtering, error correction, and chimera detection(Benjamin J Callahan, 2016). Amplicon sequencing units from DADA2 were assigned taxonomic identifiers with respect to Green Genes release 13_08 using the QIIME 2 q2-featureclassifier(Nicholas A. Bokulich, 2018). Alpha diversity with respect to: Faith PD whole tree, Evenness (Shannon) index, and observed species number metrics; was estimated using QIIME 2 at a rarefaction depth of 5,000 sequences per subsample. Beta diversity estimates were calculated within QIIME 2 using weighted and unweighted Unifrac distances as well as Bray-Curtis dissimilarity between samples at a subsampling depth of 5,000. Results were summarized, visualized through principal coordinate analysis, and significance was estimated as implemented in QIIME 2. Significance of differential abundance was estimated using ANCOM as implemented in QIIME 2 (Lin and Peddada, 2020).

### Fecal metabolite analysis by LC-MS

#### Sample preparation for LC-MS

20-30 mg of feces collected from C57BL/6J mice fed with 0.86% control diet or 0.172% methionine restricted diet were extracted in cold 80% methanol/20% water for 15 minutes at −80 °C. Extracts were then transferred into microcentrifuge tubes and centrifuged at 14000 rpm for 10 minutes. The resulting supernatants were transferred to new tubes and dried in a vacuum concentrator at room temperature. The dry pellets were reconstituted into 30 μl sample solvent (water:methanol:acetonitrile, 2:1:1, v/v) and 3 μl was further analyzed by liquid chromatography-mass spectrometry (LC-MS).

#### LC-MS method

Ultimate 3000 UHPLC (Dionex) is coupled to Q Exactive Plus-Mass spectrometer (QE-MS, Thermo Scientific) for metabolite profiling. A hydrophilic interaction chromatography method (HILIC) employing an Xbridge amide column (100 x 2.1 mm i.d., 3.5 μm; Waters) is used for polar metabolite separation. Detailed LC method was described previously (Liu et al., 2014) except that mobile phase A was replaced with water containing 5 mM ammonium acetate (pH 6.8). The QE-MS is equipped with a HESI probe with related parameters set as below: heater temperature, 120 °C; sheath gas, 30; auxiliary gas, 10; sweep gas, 3; spray voltage, 3.0 kV for the positive mode and 2.5 kV for the negative mode; capillary temperature, 320 °C; S-lens, 55; A scan range (m/z) of 70 to 900 was used in positive mode from 1.31 to 12.5 minutes. For negative mode, a scan range of 70 to 900 was used from 1.31 to 6.6 minutes and then 100 to 1,000 from 6.61 to 12.5 minutes; resolution: 70000; automated gain control (AGC), 3 × 106 ions. Customized mass calibration was performed before data acquisition. Metabolomics data analysis-LC-MS peak extraction and integration were performed using Sieve 2.2 (Thermo Scientific). The peak area was used to represent the relative abundance of each metabolite in different samples. Missing values were handled as described in (Liu et al., 2014). MetaboAnalyst package was used for the analysis of the metabolomics data (Chong et al., 2018).

### Lead sulfide assay of fecal H_2_S production

H2S production of mouse fecal samples was detected by the lead sulfide method described in Hines et al. (Hine and Mitchell, 2017). Fecal samples were collected into 1.5 ml tubes and weighed. PBS was added to each tube and samples were homogenized by vortexing and pipetting. An equal amount of fecal content from each sample was then suspended in PBS supplemented with 10 mM cysteine and 1 mM pyridoxal-5-phosphate. 150 μl of each sample was plated in 96-well plates. A piece of 703 style Whatman filter paper (VWR), soaked in 20 mM lead acetate (Sigma) and dried, was placed over the plate wells and covered with the plate lid with a heavy object on the top. The plate was incubated at 37 °C for 18 hours. The formation of lead sulfide indicated by the dark circles on the filter paper was recorded with a digital camera. The relative intensities of lead sulfide were quantified using Fiji 2.0.

### In vitro PBMC activation analysis by fecal bacteria

To test whether fecal microbes from C57BL/6J mice fed with different methionine diets have distinct abilities to activity T cells, whole blood collected from naive C57BL/6J mice fed with chow diet was treated with 1x RBC Lysis Buffer (Invitrogen, 00-4300-54) for 15 minutes to lyse red blood cells. The resulting peripheral blood mononuclear cells (PBMC) were cultured in RPMI1640 + 10% FBS in a 12-well plate at 1×10^6^ cells/well overnight. The next day, 5 fresh fecal pellets were collected from C57BL/6J mice fed with either control diet or MR diet, washed with 1x PBS for 3 times. The washed fecal microbes were then added to the cultured PBMCs with the ratio of bacteria: cells =1:1000 and incubated for additional12 hours. The same amount of PBMCs incubated with CD3/CD28 beads and PMA/ionomycin (Biolegend, Cat. # 423301) were used as a positive control. PBMCs without any treatment were used as a negative control. The fraction of IFN*γ*^+^CD8^+^ T cells were analyzed by flow cytometry.

### Quantification statistical analysis

Values are expressed as mean ± standard error of the mean (s.e.m.) from at least four biological replicates, unless otherwise indicated in the figure legend. Significant differences between the means were analyzed by the two-tailed and unpaired student’s test, and differences were considered significant at p<0.05. Data were analyzed using Prism Software 8.0 (GraphPad).

## Data availability

The RNA-seq dataset has been deposited to Gene Expression Omnibus under the accession number GSE165993.

The scRNA-seq dataset has been deposited to Gene Expression Omnibus under the accession number GSE181220.

Additional information about DEGs of RNA-seq data and IPA results is included in Table S1 and S2.

The 16S rRNA amplicon sequencing results are in Table S3.

The scRNA-seq results on immune cell clusters and T cell populations are in Table S4, S5, and S6.

The fecal metabolites data are available in Table S7.

All primers and antibodies used in the study are listed in Table S8 and S9.

## Acknowledgements

We thank Drs. Paul Wade and Michael Fessler at the NIEHS for critical reading of the manuscript, and Dr. Yisong Wan from UNC-Chapel Hill for support in FACS data analysis and critical reading of the manuscript. We also thank NIEHS Comparative Medicine Branch for support with animal experiments; NIEHS Epigenomics Core Facility for performance of RNA-seq and single-cell RNA-seq experiments; and NIEHS Flow Cytometry Center for assist on FACS analysis. This research was supported by the Intramural Research Program of National Institute of Environmental Health Sciences of the NIH to X. L. (Z01 ES102205).

## Author contributions

M. J. designed the study, performed experiments, analyzed data, and wrote the manuscript. Xiaojiang Xu and J.-L. L. analyzed RNA-seq and scRNA-seq data. Q. X. analyzed fecal bacteria abundance and fecal H_2_S production. Xin Xu performed small intestinal immune cell scRNA-seq. M. A. A.-P. performed fecal 16S rRNA amplicon sequencing and analyzed the results. J. L. and J. W. L. performed fecal metabolomic analysis by LC-MS. I. S. analyzed fecal metabolites and 16S rRNA gene amplicon sequencing results. M.J., I.S. and X.W. analyzed *in vitro* T cell activation by fecal microbes. X. L. guided, designed, and coordinated the study, analyzed data, and wrote the manuscript. All authors critically reviewed the manuscript.

## Competing interests

Authors declare no competing interests.

## Notes

https://www.ncbi.nlm.nih.gov/geo/query/acc.cgi?acc=GSE181220

https://www.ncbi.nlm.nih.gov/geo/query/acc.cgi?acc=GSE165993

### Competing Interest Statement

The authors have declared no competing interest.

### Summary of Updates

The last name of the first author was corrected.

