## Supplementary figures for "Dietary methionine restriction impairs anti-tumor immunity through gut microbiota"

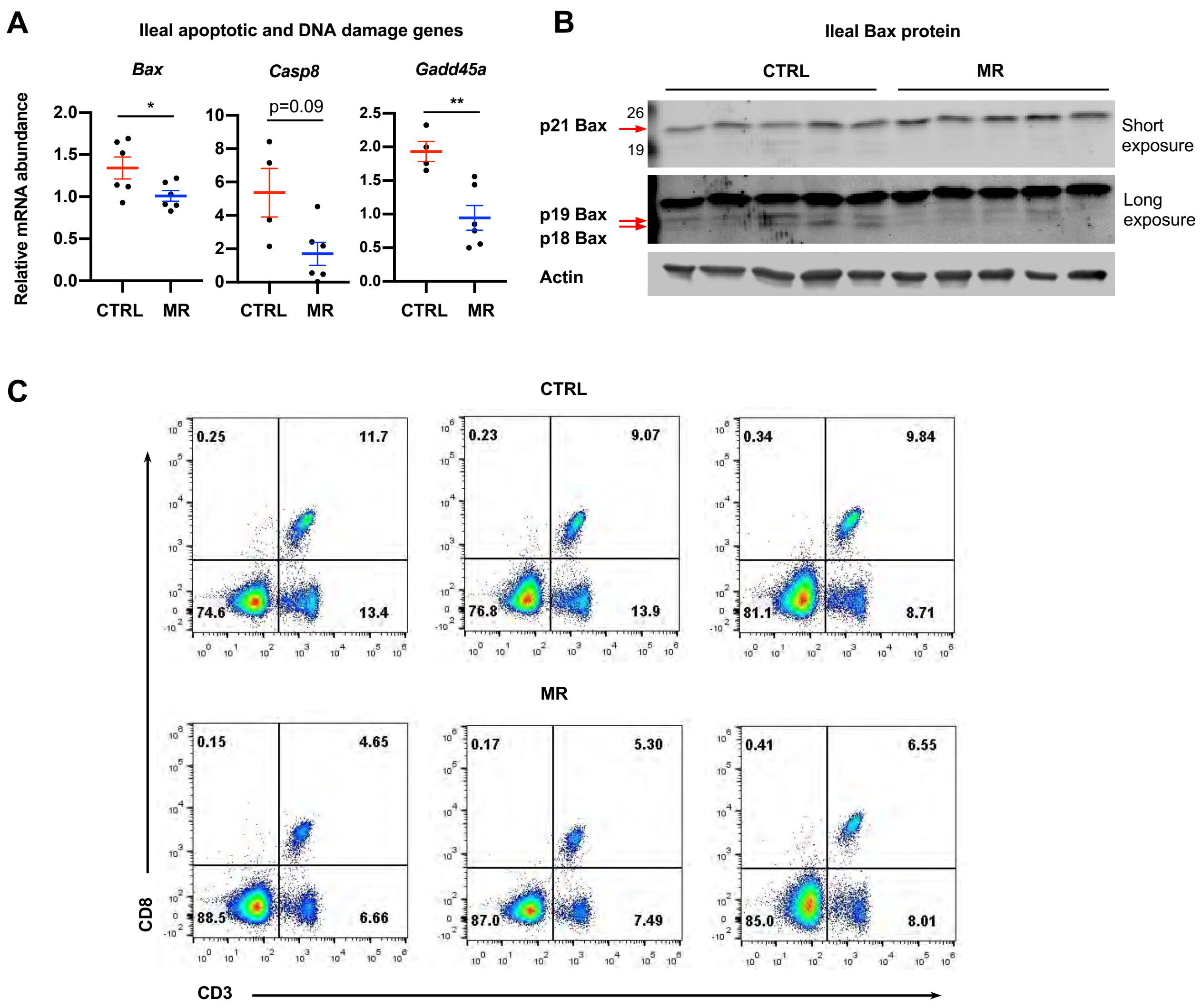


**Figure S1. Dietary methionine restriction reduces apoptosis and stress resistance in *Apc*$^{min+/-}$ mice.**

(A) Dietary methionine restriction-induced tumor growth is associated with reduced expression of apoptotic and DNA damage response genes in the ileum. The expression of indicated apoptotic genes was analyzed by qPCR (n=4-6 mice on CTRL diet; 6 mice on MR diet). (B) Dietary methionine restriction increases the cleavage of Bax in the ileum of *Apc*$^{min+/-}$ mice. The ileum sections from mice fed with CTRL diet and MR diet were immunoblotted with an anti-Bax antibody. (C) Dietary methionine restriction reduces the fraction of CD3$^{+}$ T cells in the blood. The abundance of CD3$^{+}$ T cells in the blood was analyzed by flow cytometry as described in Methods. Representative flow cytometry plots are shown.

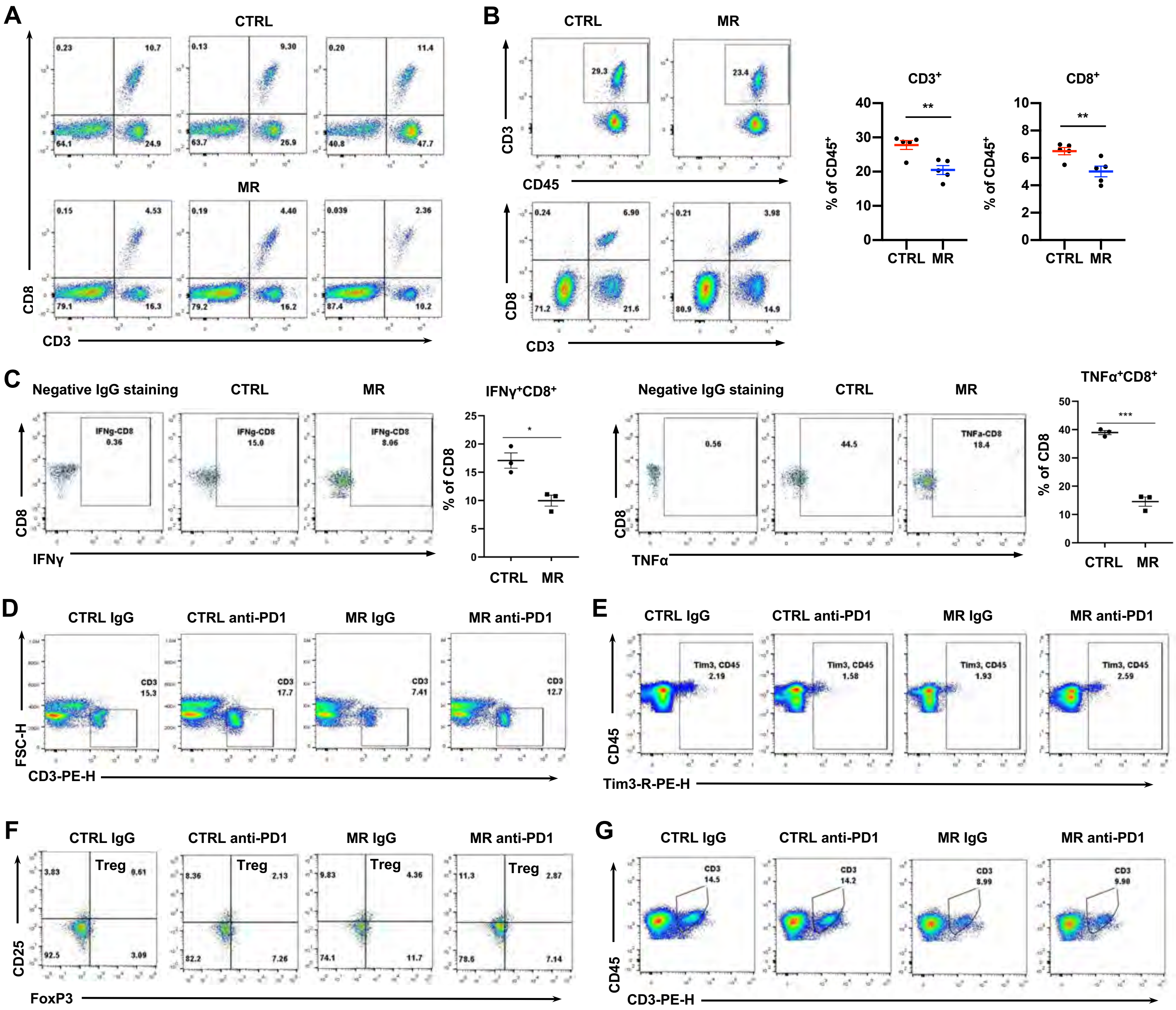


**Figure S2. Dietary methionine restriction suppresses T cell activation and dampens tumor response to anti-tumor immunotherapy in Balb/c mice.**

(A) Dietary methionine restriction reduces blood CD8$^+$CD3$^+$ T cells in Balb/c mice after two months. Representative flow cytometry plots are shown. (B) Dietary methionine restriction reduces blood CD3$^+$ and CD8$^+$ T cells in Balb/c mice after five months (n=5 mice/group, **p<0.01). Representative flow cytometry plots are shown. (C) ) Dietary methionine restriction reduces blood IFNγ$^+$ and TNFα$^+$ CD8$^+$ T cells in Balb/c mice after five months (n=5 mice/group, *p<0.05, ***p<0.001). Representative flow cytometry plots are shown. (D) Dietary methionine restriction reduced circulating CD3+ T cells. Representative flow cytometry plots of CD3$^+$ T cells in the blood from indicated treatment groups. (E) Dietary methionine restriction increases circulating Tim3$^+$ T cells Balb/c mice. Representative flow cytometry plots of Tim3$^+$ CD4$^+$ T cells in the blood from indicated treatment groups. (F) Dietary methionine restriction increases circulating Treg cells Balb/c mice. Representative flow cytometry plots of CD25$^+$ FoxP3$^+$ CD4$^+$ Treg cells in the blood from indicated treatment groups. (G) Dietary methionine restriction also reduced intratumoral CD3$^+$ T cells after the anti-PD1 antibody treatment. Representative flow cytometry plots of CD3$^+$ T cells in the tumors from indicated treatment groups

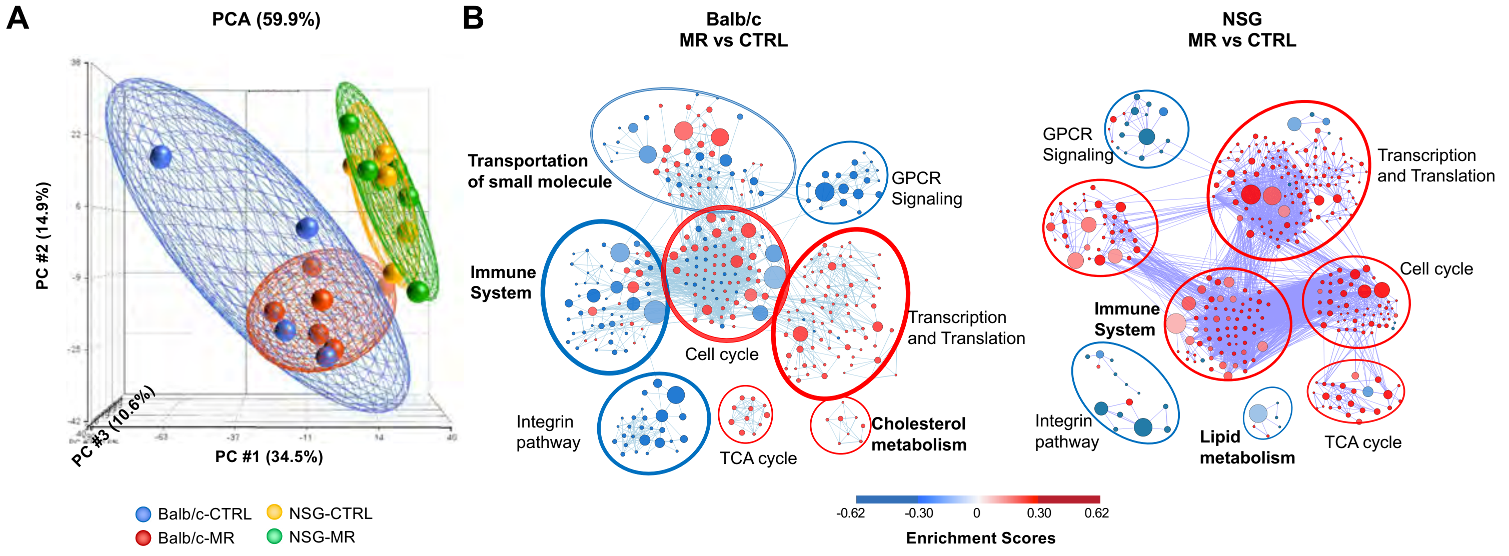


**Figure S3. Dietary methionine restriction represses immune activation in immunocompetent mice.**

(A) Dietary methionine exerts distinct impacts on the transcriptomic profiles of allografted CT26.WT tumors in Balb/c mice or NSG mice. The transcriptomes of allograft CT26.WT tumors collected from indicated mice were analyzed by RNA-seq as described (n=6 tumors in each group). (B) GSEA pathway enrichment maps for methionine restriction-induced gene sets in allografted CT26/WT tumors collected in Balb/c mice or NSG mice. Enrichment maps were generated using Cytoscape. For each node, size represents “size of gene set”, the density of color represents “Enrichment score.

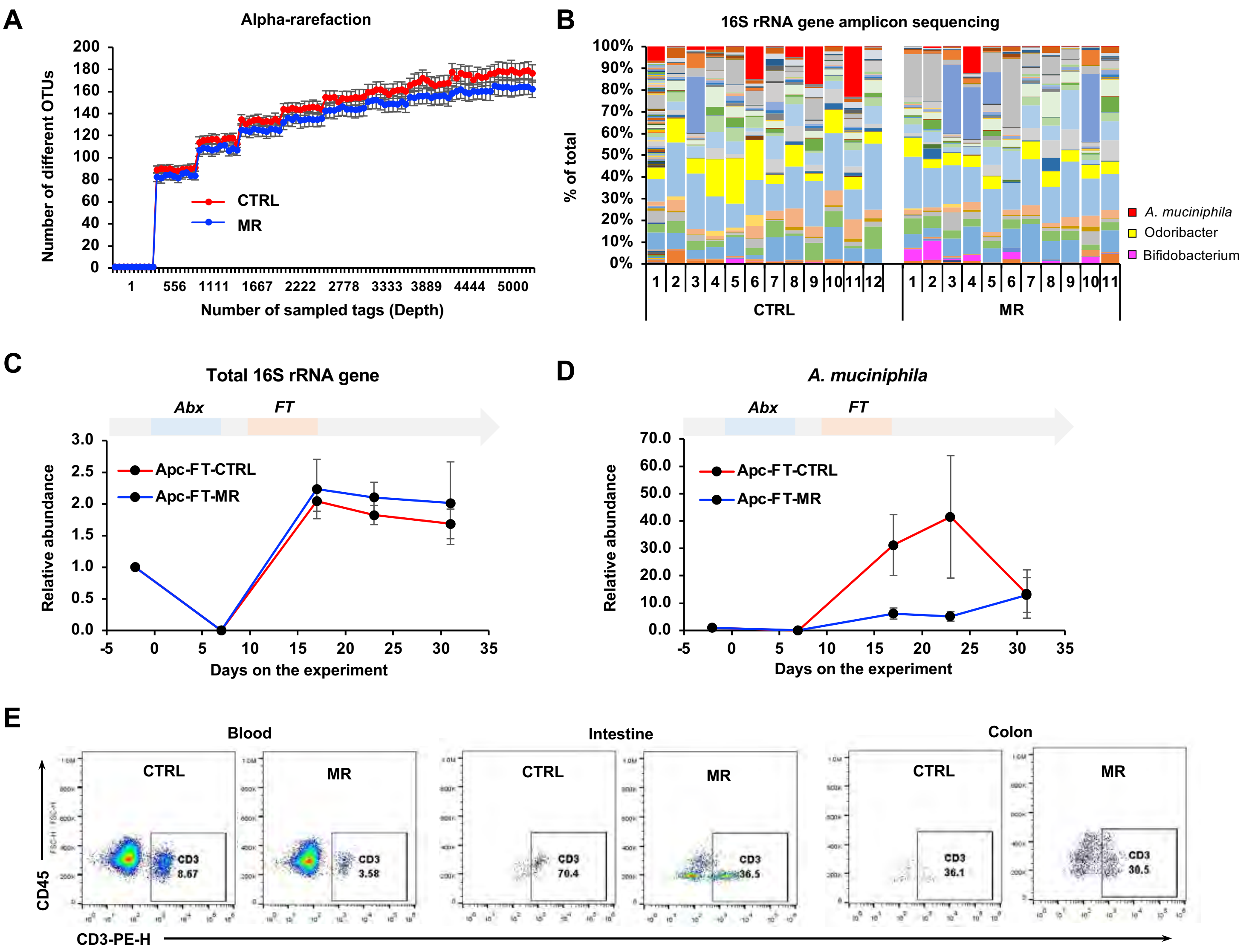


**Figure S4. Dietary methionine restriction promotes intestinal cancer progression and suppresses anti-tumor immunity through gut microbiota in immunocompetent mice.**

(A) Dietary methionine restriction slightly reduces the diversity of fecal bacteria in *Apc*$^{min+/-}$ mice. Stool DNA samples were analyzed for microbiome using 16S rRNA gene amplicon sequencing as described in Methods. Alpha diversity with respect to: Faith PD whole tree, Evenness (Shannon) index, and observed species number metrics; was estimated using QIIME 2 at a rarefaction depth of 5,000 sequences per subsample. (B) Dietary methionine restriction alters the composition of fecal bacteria in *Apc*$^{min+/-}$ mice. The fecal abundance of *A. muciniphila*, Odoribacter, and Bifidobacterium are highlighted. (C-D) Relative fecal abundance of total 16S rRNA gene and *A. muciniphila* in recipient *Apc*$^{min+/-}$ mice during antibiotic depletion of gut microbiota and fecal transplantation. The recipient *Apc*$^{min+/-}$ mice were treated with antibiotics (Abx) and fecal transplanted (FT) as described in Methods. Their feces were collected at indicated days to confirm the success of gut bacteria depletion and fecal transplantation by qPCR (n=5 FT-CTRL and 7 FT-MR). (E) *Apc*$^{min+/-}$ mice transplanted with feces from MR diet fed C57BL/6J mice have reduced abundance of CD3$^{+}$ T cells in the blood and small intestine. Representative flow cytometry plots are shown.

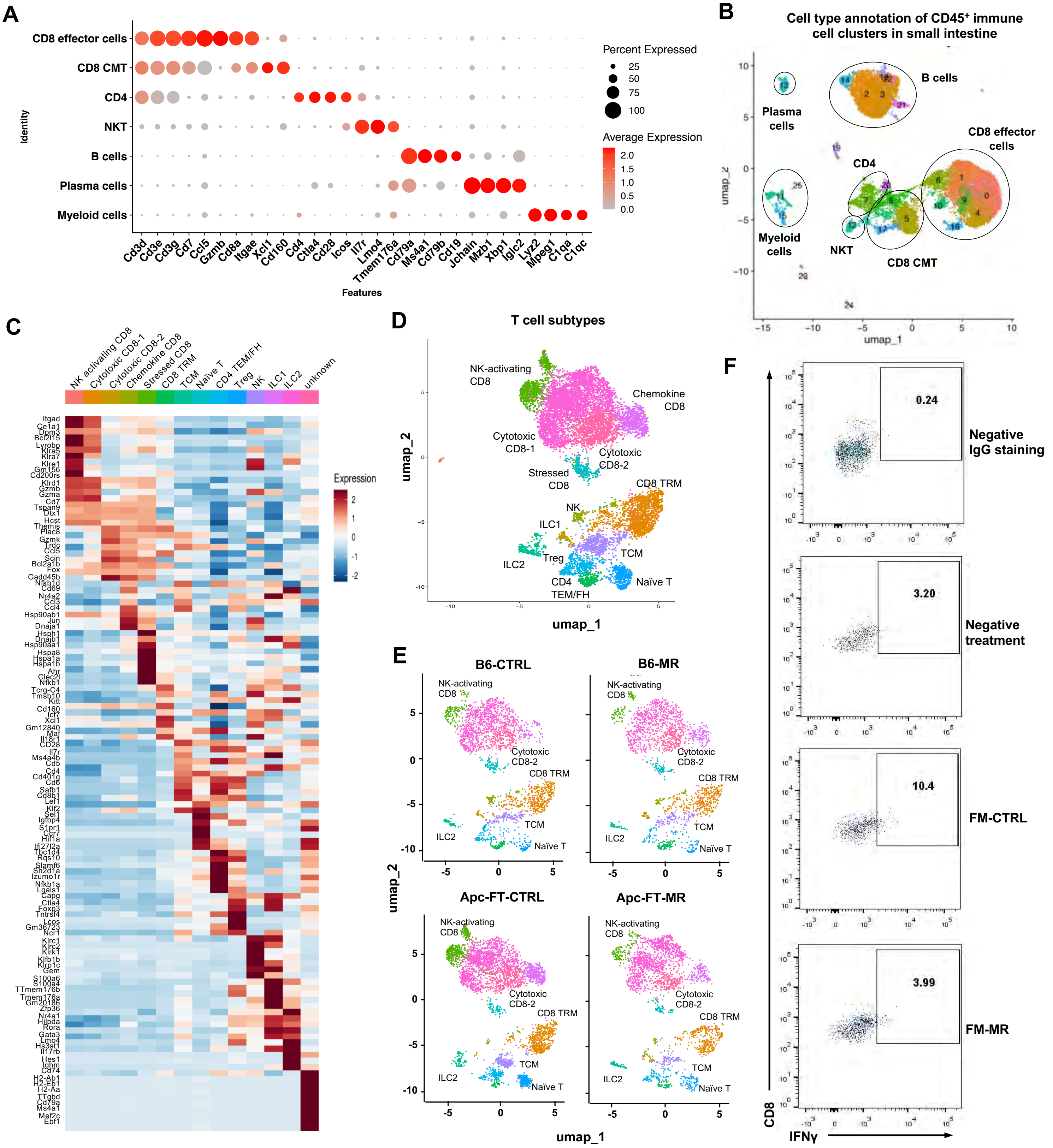


**Figure S5. scRNA-seq analysis of CD45+ immune cells from the small intestine of diet-fed B6 donor or fecal transplanted *Apc$^{min+/-}$* mice.**

(A) Dot plot showing the expression of marker genes in different function groups of total CD45$^{+}$ immune cells. Cell population annotation of CD45$^{+}$ immune cells from CTRL or MR fed B6 donor mice and from *Apc$^{min+/-}$* mice transplanted with fecal solution from CTRL or MR B6 mice. (B) Total CD45$^{+}$ immune cells from the whole small intestine of indicated mice were analyzed by scRNA-seq as described in Methods. (C) Heat map showing the expression of marker genes in different sub-groups of T cells. (D) Sub T cell group annotation. (E) Dietary methionine restriction and MR diet-trained fecal microbiota reduces the fraction of several subgroups of intestinal CD8$^{+}$ T cells. (F) Representative FACS plots for in vitro activation of T cells by fecal microbes from control diet or MR diet fed B6 mice.

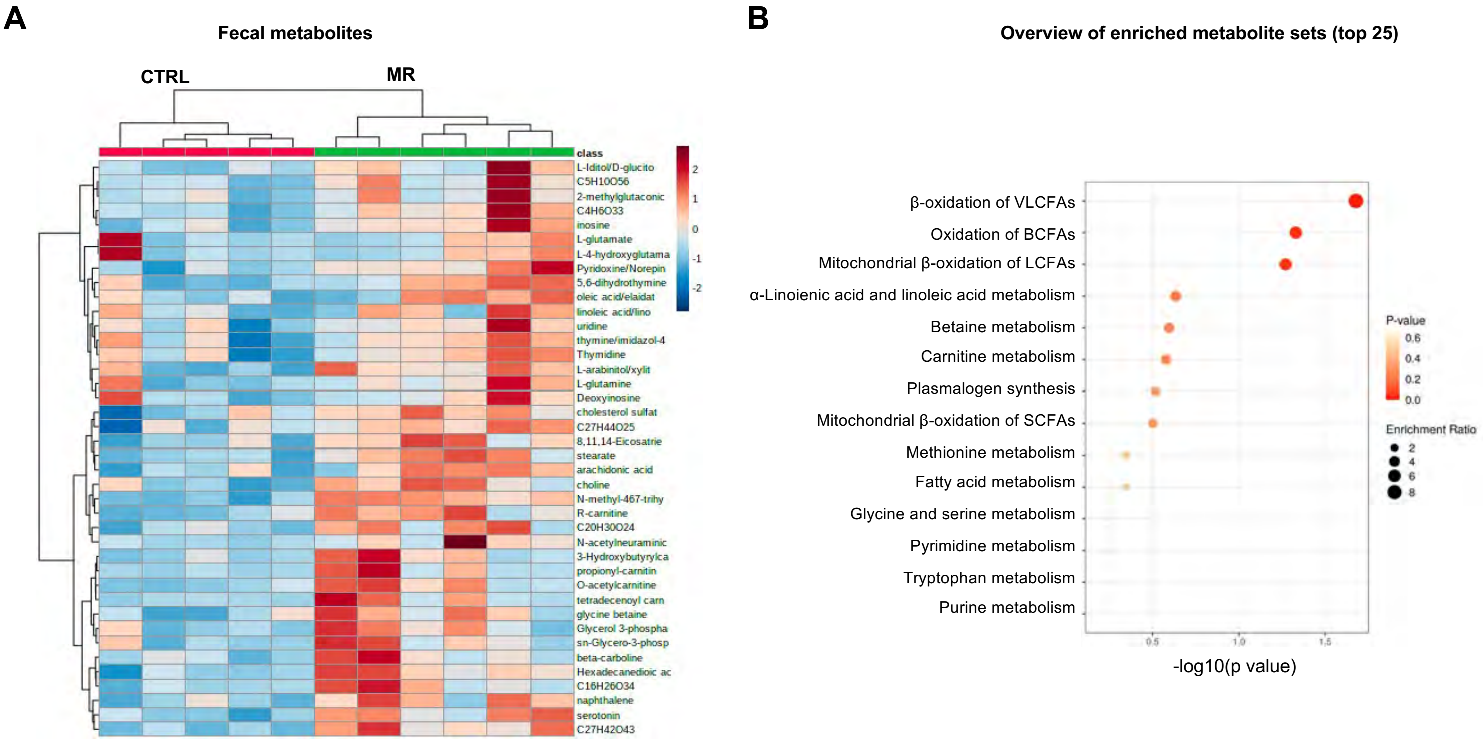


**Figure S6. Dietary methionine restriction impacts fecal metabolites.**

(A) C57BL/6J (B6) mice fed with different methionine diets have different fecal metabolites. Fecal metabolites from B6 fed with CTRL diet or MR diet were analyzed by LC-MS based metabolomics analysis. A cluster analysis was performed using all significantly ($p<0.05$, $|FC|>1.5$) changed metabolites in MR group vs CTRL group. (B) Dietary methionine restriction increases metabolites involved in fatty acid oxidation in feces of B6 mice. 19 upregulated fecal metabolites in MR fed mice were subjected for enrichment analysis using reference metabolome and SMPDB database in Metaboanalyst 5.0.

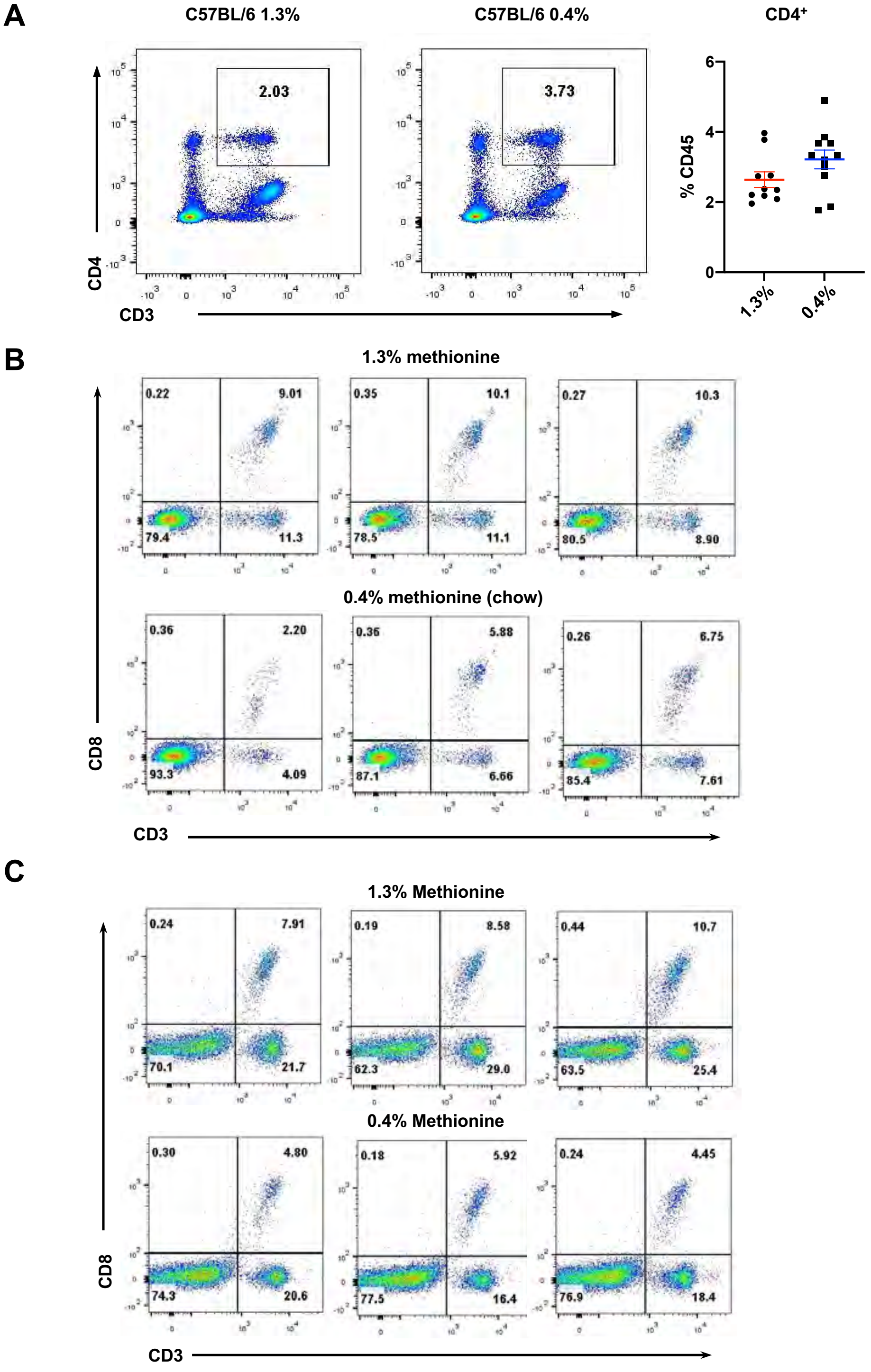


**Figure S7. Dietary supplementation of methionine increases anti-tumor immunity.**

(A) Dietary methionine supplementation does not affect circulating CD4+ T cells in C57BL/6J mice. The fractions of CD4+ T cells in the peripheral blood of mice in (A) were analyzed by flow cytometry (values are expressed as mean +/- s.e.m., n=8 mice on 1.3% diet and 11 mice on 0.4% diet). (B) Dietary methionine supplementation increases the fraction of $CD3^+$ T cells in the blood in *Apc*$^{min+/-}$ mice. Representative flow cytometry plots are shown. (C) Dietary methionine supplementation increases blood $CD8^+CD3^+$ T cells in immunocompetent Balb/c mice after two months. Representative flow cytometry plots are shown.
